# ECM Signatures Reveal Quiescent Stem Cell Diversity in the Colonic Niche

**DOI:** 10.1101/2025.01.13.632868

**Authors:** Séamus E. Hickey, Massimo Andreatta, Christina Enright, Emmanuel Boucrot, Patrick Kiely, Siobhán B. Cashman, Saintiago J. Carmona, Kieran McGourty

## Abstract

Colonic stem cells have a key role in the continuous regeneration of a healthy intestinal epithelium. Despite considerable advances in single-cell omics technologies, the transcriptional heterogeneity of rare cell types such as colonic stem cells, as well as their functional states and niche-specific behaviors, remain poorly characterised. In this study, we leverage a meta-analysis of scRNA-seq and spatial transcriptomic datasets to comprehensively map the heterogeneity of colonic stem cells. We identify multiple, previously underappreciated stem cell states, including distinct quiescent subtypes marked by *CDKN1A* (P21), *CDKN1B* (P27), and *CDKN1C* (P57), proliferative populations defined by *MKI67* (Ki67) and *LRIG1*, and a lineage-committed intermediate subpopulation expressing *MUC2*. Strikingly, we find that these states can be robustly identified solely by their extracellular matrix (ECM) gene expression signatures, revealing ECM composition as a critical determinant of stem cell identity. Notably, LAMA1 expression is highly specific to the P57+ quiescent population, linking laminin-mediated microenvironments to the active maintenance of deep quiescence, consistent with our recent findings associating LAMA1 with quiescent cell survival. By applying these ECM gene signatures, we delineate discrete “micro-niches” of quiescent stem cells in healthy tissue and provide evidence that analogous states persist in the colorectal cancer (CRC) environment. Extending our approach to an unrelated tissue, the pancreas, we detect parallel quiescent cell subtypes, illustrating the broader applicability of ECM-based signatures. Taken together, our findings redefine the concept of stem cell heterogeneity in the colon, establish ECM-driven gene signatures as a powerful tool for characterizing stem cell states, and offer new perspectives on the niche-dependent regulation of both healthy and cancerous stem cell populations.

## Introduction

Stem cells are classically defined as having the ability to indefinitely self renew and to give rise to daughter cells that can differentiate into various cell types^1^. These properties are central to the remarkable regenerative capabilities of organs in adult tissues, with adult stem cells acting as key players in tissue maintenance and repair^2^. They are multipotent and produce limited numbers of cell types associated to the tissue of origin, unlike embryonic stem cells. They are also important for homeostatic tissue maintenance, wound healing, barrier function, innate immune responses to infection and the aging process.^3^.

These adult stem cells reside within defined anatomical regions known as the stem cell niche. Within these niches, stem cells can exist in a spectrum of functional states, including deep quiescence, primed quiescence, proliferative states, and others displaying hallmarks of differentiation^2^. While it is clear that these varied states contribute to tissue integrity and regenerative potential, a complete understanding of the full range of possible states within the niche remains elusive. Moreover, the niche determinants that orchestrate transitions between these states are only partly understood.

To further increase our knowledge of these regulatory mechanisms, it is important to consider the stem cell niche environment, which comprises both cellular and non-cellular components. The major non-cellular element is the extracellular matrix (ECM), which offers biophysical and biochemical cues that influence stem cell behavior. A complex microenvironment that comprises gradients of soluble molecules (e.g., growth factors, hormones), insoluble molecules (e.g., ECM components), and mechanical features directs controlled activation and hierarchical differentiation^4^. A growing body of evidence acknowledges the ECM as a key regulator of stem cell quiescence; an inactive, protective state that preserves stem cell populations over time^5,6^. In various tissues, quiescent stem cells are spatially distinguished from active stem cells, indicating that distinct niche microenvironments maintain these states^7^.

Importantly, quiescence is not a singular state. It encompasses heterogeneous sub-populations, including deeply dormant G0 cells, preparatory G-alert cells, and other potential transcriptional phases^8,3^. Whether these represent discrete states with unique molecular programs, or merely snapshots along a continuum, remains an open question. Still, certain core molecular signatures are consistently observed. Cyclin-dependent kinase (CDK) inhibitors such as *CDKN1A/P21, CDKN1B/P27*, and *CDKN1C/P57* are frequently identified in quiescent cell populations across multiple tissues^2,9,10,11^, blocking cell cycle progression and preventing unwarranted proliferation. Tissue-specific examples include *P57* governing quiescence in the colonic epithelium^12^, *P21* maintaining quiescence in the hair follicle stem cell niche^9^, and *P27* supporting quiescence in the hippocampus stem cell niche^10^. Yet, it remains uncertain how universal these regulators are across diverse tissues, cell states and niches.

Recent work from our group highlights the complexity of ECM-mediated quiescence regulation. We found that laminin 111 can induce a reversible exit from the cell cycle into G0 and promote long-term survival mediated by sustained MAPK/ERK signaling. Activated by laminin-binding Integrin *α*3*β*1 through the Rap1-BRAF-MEK pathway and anchored by IQGAP1 in the cytoplasm, elevated pERK prevents nuclear entry and thus blocks proliferation, while simultaneously ensuring cell survival by suppressing pro-apoptotic proteins like BAD. Moreover, we confirmed that cytoplasmic pERK remains elevated in *LGR5+* stem cells within Laminin *α*1-positive niches in the porcine intestine^13^, directly linking a specific ECM component to the active maintenance of quiescence.

The colonic intestinal epithelium exemplifies a structured system where such dynamics can be closely examined. Composed of crypts of Lieberkühn that harbor stem cells at their base, the epithelium undergoes continuous regeneration, renewing every four to five days^14,15^. Stem cells in the crypt-base zone drive a hierarchical lineage progression through transient amplifying (TA) zones before reaching differentiated compartments^16^. Although often depicted as cycling, many of these stem cells remain mostly quiescent, and evidence points to multiple quiescent states within these niches^2^. The ability of these cell populations to transition between quiescence and the cell cycle under homeostatic or injured conditions is critical for tissue integrity.

However, even well-studied markers of quiescence have proven contentious. For example, *LRIG1* was proposed as a marker of quiescent intestinal stem cells at the +4 position along the crypt axis. Yet, subsequent studies found a significant population of *LRIG1* in *LGR5*^*+*^ cells found to be cycling as marked by *Ki67*^*+*17^, raising doubts about its specificity as a quiescence marker and suggesting that certain “canonical” quiescent markers may reflect active states instead.

The importance of accurately defining stem cell states extends beyond healthy tissues. In diseased states, such as colorectal cancer (CRC), both proliferation and quiescent cycles influence oncogenic processes, including tumorigenesis, progression, treatment resistance, and recurrence^18^. Cancer stem cells (CSCs), like their healthy counterparts, reside in niche environments where ECM composition can dictate whether they remain dormant or actively divide. In CRC, CSC quiescence marked by *LGR5*^*+*^/*P27*^*+*^ cells was shown to be regulated through the ECM protein *COL17A1* via FAK-YAP activation^19^. This is particularly significant since quiescent CSCs are implicated in therapeutic resistance and tumor relapse^20,21^. Whether ECM-mediated quiescent regulation in healthy tissue parallels that in cancer remains an open question.

A major challenge in resolving these complexities has been the rarity of stem cells and the difficulty in distinguishing their signals from those of surrounding cells^22^. However, recent technological advancements in single-cell RNA sequencing (scRNA-seq), spatial transcriptomics (ST), and spatial proteomics (SP) have made it possible to examine cell states and behaviors at unprecedented resolution^23^. These tools enable the dissection of cellular heterogeneity, previously masked by bulk tissue analyses, and open new avenues for identifying distinct stem cell states and their niche determinants.

Accordingly, this study leverages these technologies to provide a more complete understanding of the various states stem cells can adopt in the healthy colon. We further investigate the interdependence of ECM components in regulating these states. Finally, we examine whether similar states and ECM dependencies occur in CSCs within colonic tumors, exploring the heterogeneity of gene expression and niche ECM dysregulation. Using these data, we generated ECM signatures of stem cell quiescence in both healthy and diseased tissues and validated their capacity to identify dormant sub-populations in spatial transcriptomic sections of colonic tumors. Lastly, we tested the broader applicability of these quiescent ECM signatures by identifying quiescent stem cell sub-populations (stellate cells) in healthy pancreatic tissue, a distinct anatomical region from which the original signatures were derived.

To our knowledge, this represents the most comprehensive description of the various states of stem cells within their niches to date. Moreover, it reveals the intricate molecular underpinnings linked to ECM composition that help define, maintain, and potentially manipulate these states.

## Results

### Single-cell RNA sequencing distinguishes stem cells from differentiated epithelial cells

We first examined our ability to successfully isolate stem cells from a mixed epithelial single cell dataset comprising cells sequenced from 38 donors and 142,113 epithelial cells generated as part of the Human Gut Atlas project. We chose this dataset as a model of the epithelial stem cell lineage in the colon (Fig 1. S1a) and to challenge the specificity of our stem cell isolation approaches amongst cells with related characteristics. We subset our populations based on the expression of epithelial cell lineage marker EPCAM in conjunction with either LGR5, to identify stem cells, or CEACAM7 to identify differentiated epithelium and yielding 12,282 cells. After applying dimensionality reduction, six distinct clusters emerged in the reduced feature space(Fig 1. S1b). These cells were then inspected for their expression of *LGR5* and *CEACAM7* indicating clear separation between clusters positive for each, allowing for confident annotation of stem cells and differentiated epithelial (Diff Ep) respectively (Fig 1. S1c,d).

**Figure 1.**
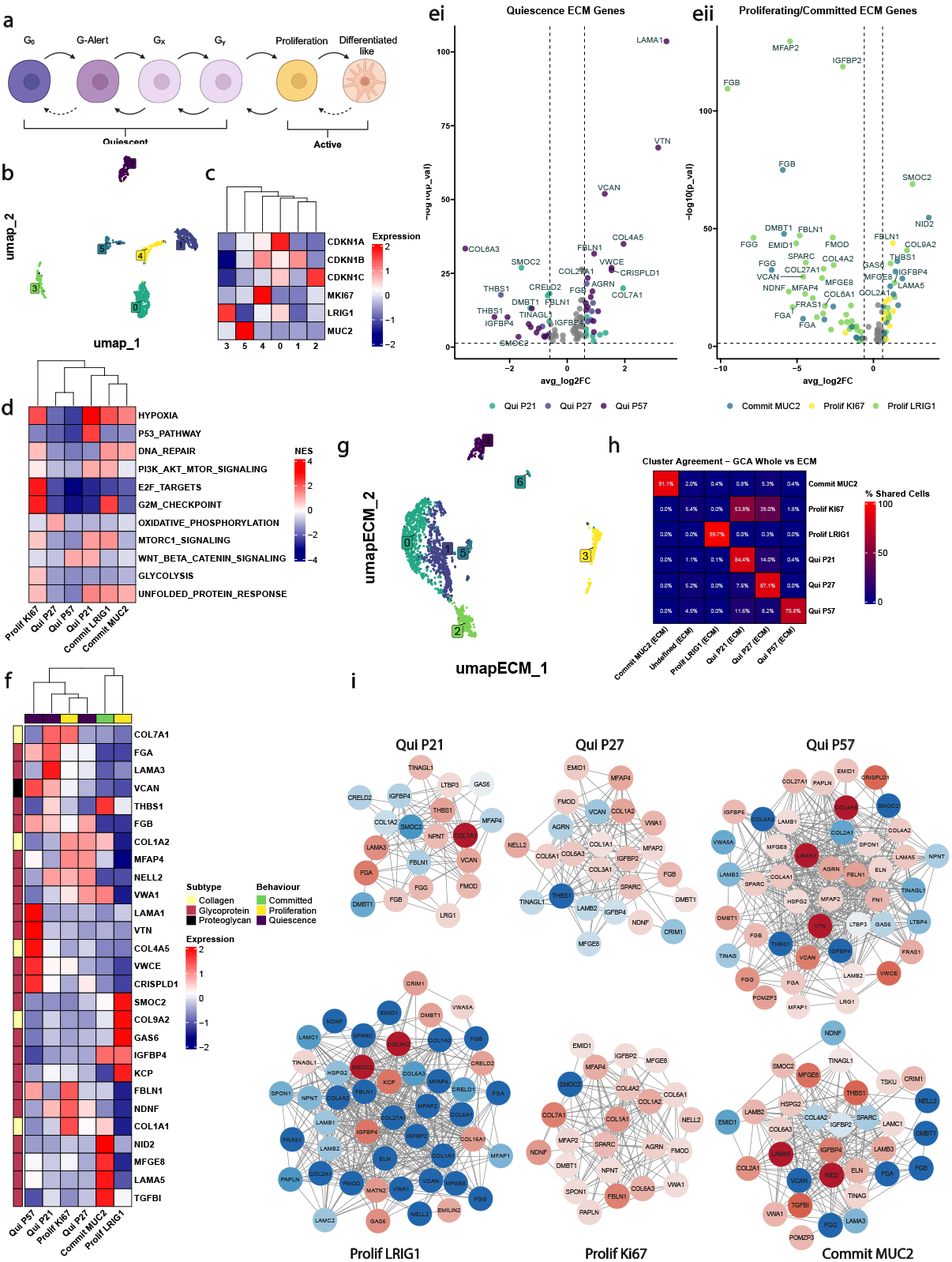
Colonic stem cell heterogeneity from scRNA-seq meta-analysis. **(a)** Progression of somatic stem cell through varying stages of activation and differentiated like states. **(b)** UMAP showing six clusters of transcriptionally unique epithelial stem cells. **(c)** Normalized, rescaled expression for key marker genes in six stem cell clusters. **(d)** Clustered heatmap of Pathway Inference Analysis performed on annotated clusters of stem cells against the PROGENy database gene sets. **(ei)** Volcano plot of ECM differentially expressed genes (DEGs) in the quiescent-like clusters, as per their Average Log 2 Fold Change. **(eii)** Volcano plot of ECM DEGs in the active-like clusters, as per their Average Log 2 Fold Change. **(f)** Clustered heatmap of the top 5 Core Matrisome ECM DEGs expressed per cluster. **(g)** UMAP depicting clusters of epithelial stem cells generated when only using ECM genes as defined by the Matrisome DB. **(h)** Agreement matrix showing the percentage of shared cell between the clusters generated using only ECM genes and the clusters generated in **Fig 1 (b)** using the whole RNA transcriptome. **(i)** Network analysis of all ECM based DEGS within each of the clusters from **Fig 1 (b)**

To further support the accuracy of our isolation approach for stem cells versus diff ep. we assessed the expression of known markers of stemness and differentiation including the original markers used to isolate these cells from their original datasets (*EPCAM/LGR5/CEACAM7*). Consistently, the annotated population of stem cells showed increased expression in stem markers *ALDH1A1, ASCL2* and *SOX9* while differentiated epithelial cells highly expressed *KRT20, CEACAM1* and *GUACA2A* (Fig 1. S1f, g). Lastly, we investigated the purity of our populations in terms of their crypt-axis score, a gene signature that measures the position of a cell from the bottom of the crypt (undifferentiated stem cells) to top of crypt (fully differentiated colonocytes)(see Methods Fig 1. S1h). These data support the accuracy of our stem cell isolation with LGR5+ stem cells and CEACAM7 positive cells aligning to the expected positions within the colonic crypt.

### Colonic stem cell niche determinants driving stem cell state heterogeneity

Next, we focussed on stem cells based on the positive expression of *EPCAM* and *LGR5* and negative expression for *CEACAM7*. Unsupercised clustering of scRNA-seq resulted in 6 transcriptionally unique populations of stem cells (Fig 1b). These 6 clusters were characterised by markers previously associated to stem cell populations of the colon, including: *CDKN1A* (P21), *CDKN1B* (P27), *CDKN1C* (P57), *MKI67* (KI67), *LRIG1* (LRIG1) and *MUC2* (MUC2) (Fig 1c). Restricted expression of these markers allowed us to further categorise each cluster by their cell state of quiescence, proliferation or commitment^2,24–26^. Accordingly, we annotated our clusters against these reporter gene patterns resulting in *Quiescent P21 (Qui P21), Quiescent P27 (Qui P27), Quiescent P57 (Qui P57), Proliferation KI67 (Prolif KI67), Proliferation LRIG1 (Prolif LRIG1)* and *Committed MUC2 (Commit MUC2)*.

While the conventional paradigms regard the adult stem cells as a single class controlled by the limited set of the canonical pathways, our results reveal a significant heterogeneity in the stem cell population. In order to study the variability of these populations in terms of pathway activation and metabolic activity, we performed pathway inference and gene set enrichment analysis (GSEA). This analysis revealed specific regulation in key cell cycle, metabolic activity and stem related pathways across the three quiescent (Qui P21, Qui P27, Qui P57), two proliferative (Prolif KI67, Prolif LRIG1) and one committed (Commit MUC2) clusters we identified. Overall, this analysis points to a hierarchy of activation in this compartment, with Qui P27, Qui P57 displaying the most deeply dormant signatures with reduced metabolic activity This more detailed characterisation, which includes the detailed annotation of the pathways for each subpopulation. Here, we explain how certain pathways like WNT, PI3K/AKT/mTOR, DNA repair, cell cycle and Glycolysis differ between the quiescent (Qui P21, Qui P27, Qui P57) and/or proliferative (Prolif KI67, Prolif LRIG1) and/or committed (Commit MUC2) stem cell states, which point toward a more complex and layered regulatory network than what has been thought (Fig 1d, Fig1. S2).

Given the cell state transitions outlined above and the extensive literature supporting and our own findings implicating the ECM as being an integral factor governing cell behaviour^27,28^, we examined whether ECM genes were part of the DEGs influencing our clusters and shortlisted them against a reference list generated from the MatrsiomeDB^29^, as displayed on a superimposed volcano plot grouping quiescent clusters and proliferative/committed clusters (Fig 1e) This examination highlighted specific ECM signatures associated with each of our cell states, raising the possibility that niche associated ECM extrinsic signalling could be regulating each state. Consistent with our groups own findings^13^, *LAMA1* was heavily regulated to a specific subset of LGR5 positive stem cells, here characterised by P57 expression, together with *COL4A5* and *COL7A1* in quiescent clusters Qui 57 and Qui 21 respectively, while *COL6A3* was downregulated. Interestingly, *SMOC2, VCAN* and *THBS1* were inversely regulated in across multiple quiescent clusters when compared to proliferating/committed groups. *THBS1* has previously identified a modulator of stem cell self-renewal and senescence through CD47^30^. SMOC2 has been identified as labelling an LGR5 sub-population, though this was reported to be enriched in *LRIG1* slow cycling clusters^31^ rather than quiescent populations as we find here (Fig 1e-f). VCAN has been associated with several activated cell processes such as proliferation, differentiation and cell adhesion amongst others^32^ and surprisingly we find it up regulated in the quiescent niche here, again highlighting some confusion in the field while substantiating the need for thorough characterisation followed here.

Following these observations, we reasoned ECM gene regulation could be the primary drivers of the six cell states of the LGR5 stem cell compartment we observed. To test this hypothesis we re-clustered our cell populations solely using ECM genes, under the assumption that cell states regulated by a specific ECM signature would similarly cluster with their associated ECM genes. This approach yielded 7 clusters (Fig 1g) and upon investigating the expression of the canonical state markers found previously we found that LRIG1, P21, P27, P57 and Muc2 displayed enriched expression in distinct clusters, with LRIG1 shared across 2 (FIG S1D). Interestingly, we did not find selective enrichment for Ki67, indicating that the transcription programs associated are not regulated by a unique ECM signatures in our dataset in the stem cell niche. Further, cluster 5 was not associated to any of the canonical state marker. Accordingly we annotated our ECM driven clustered data as *Qui P21 (ECM), Qui P27 (ECM), Qui P57 (ECM), Undefined (ECM), Prolif LRIG1 (ECM)* and *Commit MUC2 (ECM)* (Fig 1. S1d).

These data indicate that ECM alone was sufficient to drive annotation for 5 of the stem cells states we have described. To challenge the validity of this approach we calculated the percentage of shared cells from our ECM only clustering method with that based on the original whole assay, hypothesising that there should be a high percentage agreement were ECM genes the underlying determinants of the states observed. Compellingly, we observed excellent agreement across all subgroups with the exception of those cells of proliferating/*Prolif KI67* cluster (Fig 1h). Further when labelled according to the original whole assay cluster identity in this ECM UMAP space we failed to detect a distinct clustering of Ki67 positive cells. These findings suggest that though Prolif Ki67 cells represent a transcriptionally distinct cell state they are not governed by a specific ECM (Fig 1h). These results are conceptually consistent with the the postulate that quiescence and differentiation are active processes elicited from the immediate cellular environment and that the proliferative process requires a relief from those signals.

Lastly, we examined through protein interaction networks the core ECM gene sets regulated to the various states of the we described and observed highly connected networks reflecting a solution set of ECM genes that can regulate the 6 of the cell state behaviors, of stem cells in the niche and may represent functional modules to drive regulated stem cell states (Fig 1i). These modules appear to provide a central axis of the stem cell niche, influencing their respective phenotypes and as such warrant further investigation to evaluate their sufficiency and requirement in controlled *in vitro* or animal models.

### Non-negative Matrix Factorisation defines Gene MetaProgrammes that can describe stem cell heterogeneity

Having mapped the cells states of LGR5 intestinal stem cell niche, we wished to analyze these populations using an analogous but independent approach of gene discovery programs based on non-negative matrix factorisation. This holds the advantage of allowing us to interrogate our data without undergoing the standard integration, normalisation, and PCA dimensionality reduction typically seen in single-cell analysis pipelines. Furthermore, by deploying this approach over our data, individual gene programs can be assembled into meta-program units (MP)s based on their signal as determined through a calculated Jaccard index, allowing that some states may have MPs shared amongst them.

This analysis identified 7 discrete gene MPs present in the stem cell niche (Fig 2a) that can be used to describe the heterogeneity of this compartment. Importantly, multiple metaprogrammes may be activated in an individual cell and their degree of activation can be quantified using a UCell signature score. Thus, we were able to asses the level of activation of individual metaprogrammes according to the cell state clusters we initially described (Fig 2c, Fig 1. S2a,b). We further characterised these clusters with overlap gene set enrichment analysis of hallmark biological processes for each program identified pathways that provided a collective impression as to the processes active in cells (Fig 2b): MP1 and were dominated by processes of cycling cells with expressing of genes related to G2M checkpoint, mitotic spindle and E2F targets. MP2 was enriched in the metabolic process of oxidative phosphorylation. MP3 showing modulation in KRAS signalling, glycolysis, Estrogen response factors and EMT, a process commonly identified in the stem cell development and activation^33,34^. MP4 also showed enrichment in EMT along with WNT and coagulation. MP5 was heavily dominated by hypoxia and KRAS signalling, both processes associated with stem cells previously^33,35^. Lastly MP6 was heavily associated with NFKB regulation and to a lesser extent with hypoxia, P53 and apoptosis (Fig 2b,g).

**Figure 2.**
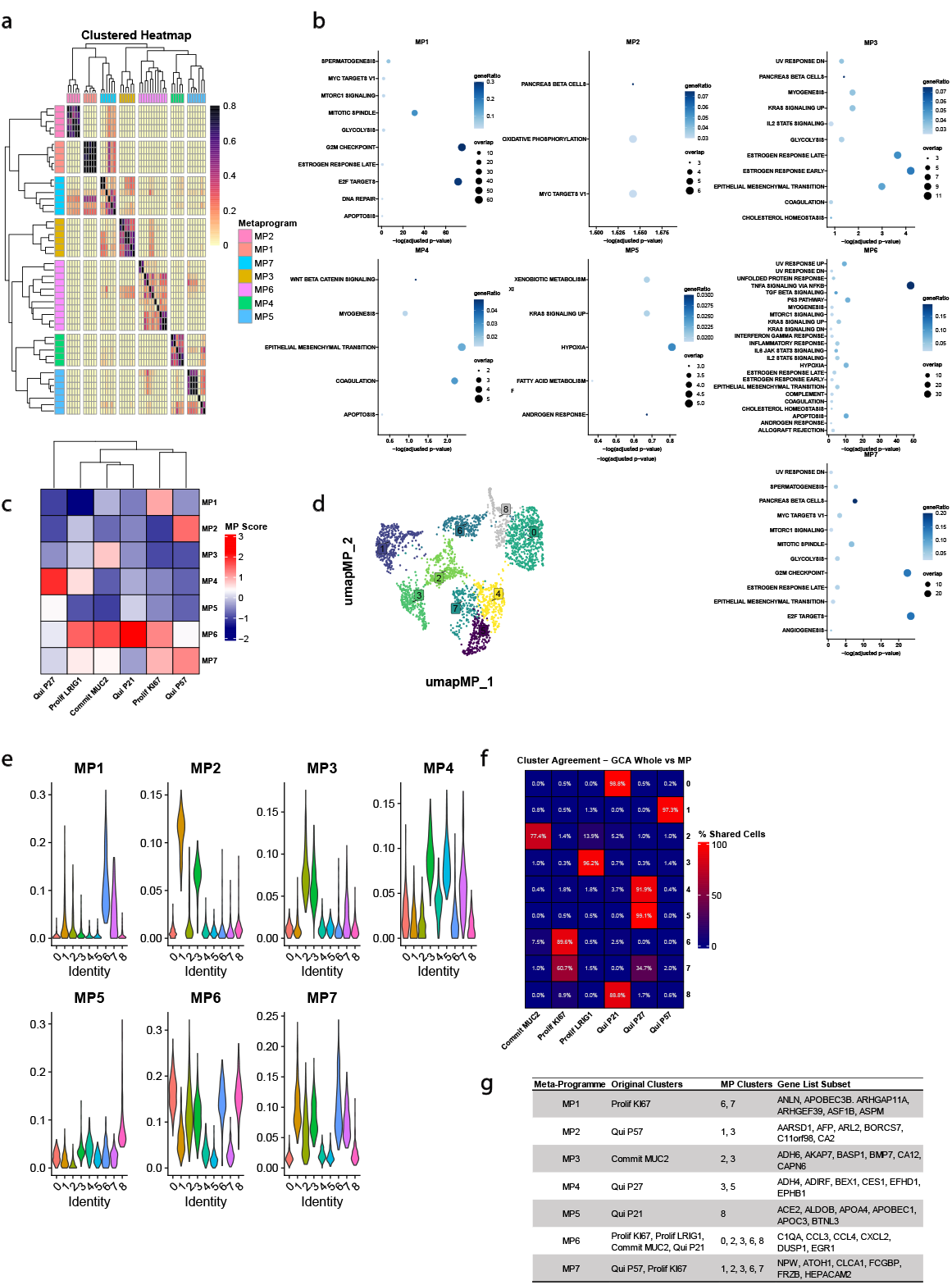
**(a)** Jaccard plot of the individual gene programmes that comprise the Meta-propgrammes (MPs). **(b)** Dotplot describing the GSEA Hallmarks that are described by each of the seven Meta-programmes. **(c)** Clustered heatmap of the relative MP signatures in the original clustering method in **Fig 1 (b). (d)** UMAP of cells clustered using the 7 dimensional space created by the relative signature scores in each cell for the MPs. **(e)** Violin plots of the expression of of each MP score in the clusters generated in **Fig 2 (d). (f)** Agreement matrix showing the percentage of shared cells between the clusters generated from this clustering method and those in the original whole transcriptome clustering in **Fig 1 (b). (g)** Summary table of the relationships between the MPs, the original whole cell clustering identities, the new MP derived clusters, and a subset of the genes that drove the original MP formation in **Fig 2 (a)**.

Though specific metaprograms were enriched in the cell states described, we wished to exploit the quantitative combinatorial evaluation inherent to this analysis and further clustered our data across the 7 dimensions of the metaprogrames (Fig 2d). We observed 9 discrete cell clusters that were variously enriched for individual MPs and were then able to asses for agreement of cell composition to the clusters we previously generated (Fig 2d-f). An advantage of clustering data in MP space is that it can provide a quantitative assessment of the degree to which the underlying biological processes of each MP contributed to various stem cell states and whether this regulation was positively or negatively enriched compared to the other states of the compartment using ranked gene set enrichment analysis. This analysis revealed significant similarities among cells clustered in MP space or cells clustered using standard methods and is discussed in more detail in SI note 2.

Remarkably, this analysis resulted in near perfect concordance between the MP space clustering and that of our initial whole transcriptome standard analysis as assessed through cell clustering (Fig 2f) or by pathway analysis (Fig 1. S2a,c). Examining intestinal stem cells through the lens of MPs can accurately recapitulate the diverse heterogeneity of these populations and allows established relationships among MPs, the original clustering method, the new MP clusters, and a subset of the genes composing the original MPs are summarised in (Fig 2g). This data further reinforces the heterogeneity we have observed in the colonic stem cell epithelium population and allows for deep understanding of the cellular processes influencing driving these state.

### ECM UCell Signature creation

After establishing our ability to define colonic stem cell heterogeneity using whole transcriptome data, ECM genes, and MPs quantitatively, we aimed to integrate these independent analyses into a more cohesive approach. To accomplish this, we identified the intersection of signatures derived from the differentially expressed genes (DEGs) within the clusters of each method, revealing consistent signals centered on ECM-associated genes (Fig 3a). This process resulted in five ECM signature gene sets that were consistently detected across all analyses: P21.Sig_UCell, P27.Sig_UCell, P57.Sig_UCell, MUC2.Sig_UCell, and LRIG1.Sig_UCell. Ki67 ECM-related signatures were variable and therefore were not brought forward here. Next, we calculated a UCell score for each cell based on each of the five remaining consensus ECM gene signatures. Evaluating the expression levels of the canonical cluster markers used earlier, we observed a strong positive correlation between the expression of each feature marker and its corresponding UCell ECM signature, particularly between the quiescence signatures and their respective markers (Fig 3b).

**Figure 3.**
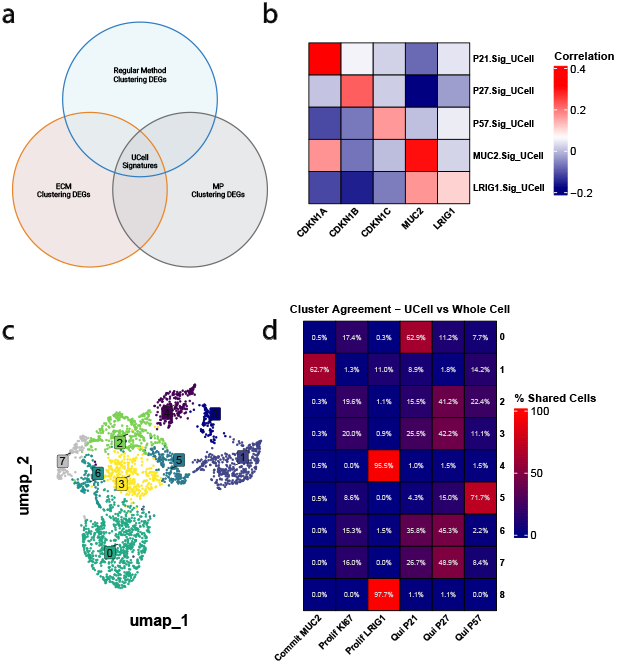
**(a)** Methodological approach that was used to generate the extracellular matrix based molecular signatures for each of the subpopulations of stem cells. **(b)** Heatmap that shows the correlation between the extracellular matrix molecular signatures that were generated to describe various stem cells states in the healthy intestine and the expression of the markers CDKN1A/P21, CDKN1B/P27, CDKN1C/P57, LRIG1 and MUC2. **(c)** UMAP showing nine clusters of stem cells after using a 5-dimensional extracellular matrix signature score against the dataset. **(d)** Agreement matrix showing the percentage of shared cells between the clusters generated using a 5-dimensional extracellular matrix signature score and the clusters generated in **Fig 1 (b)**.

Additionally, we assessed the specificity of our signatures against the various clusters we had previously identified. We found excellent agreement between the expression of ECM UCell signatures and our original clusters: Qui P21, Qui P57, Commit MUC2, and Commit LRIG1 (Fig 3. S1). Although the P27.Sig_UCell was less specific, it still showed its highest signature score in quiescent clusters (Fig 3. S1), potentially highlighting the dynamic nature of these cells as they transition through quiescent compartments.

Subsequently, we compared the signature scores with our previous clustering approaches, including ECM-driven clustering and non-negative matrix factorisation (NMF) clustering. Consistently, the UCell ECM signatures showed strong concordance with the corresponding clusters for P21, P57, MUC2, and LRIG1. However, P27 remained somewhat indistinct, and Ki67 did not exhibit a specific signature (Fig 3. S1).

Given the plasticity among the states within the stem cell niche, we hypothesised that it might be possible to cluster our data using the UCell-ECM signature scores across all five dimensions. This approach allowed us to delineate nine distinct cell populations within this five-dimensional space (Fig 3c). Utilising these data, we constructed a similarity matrix reflecting the percentage of cells shared between the original whole transcriptome clustering analysis and our UCell ECM signatures. This resulted in strong agreement for Qui P21/57, Prolif LRIG1, and Commit MUC2 clusters (Fig 3d).

As before, Qui P27 displayed some variation and was shared across clusters 2, 3, 6, and 7, with over 40% of the cells in these clusters originating from the Qui P27 cluster. Importantly, these clusters were adjacent neighbors in the UMAP space, indicating similarity, and approximately 80% of cells within each of these clusters arose from quiescent clusters Qui P21/27/57. As expected, individual UCell ECM signatures did not accurately capture the cells of the Prolif Ki67 cluster identified in the whole transcriptome analysis.

Our findings reveal a high degree of interdependence between specific ECM signatures and cell states within the LGR5 compartment. Moreover, we developed quantitative ECM signatures that can robustly identify these states, potentially enabling quantitative annotation of these behaviors in a cell-agnostic manner.

### Stem cell heterogenity in colorectal cancer epithelial stem cells

Cancer stem cells (CSCs) of colorectal cancer (CRC) are associated are associated with numerous oncogenic processes, such as cancer initiation, progression, metastasis, resistance to therapy, and recurrence, with resistance and recurrence being heavily linked to stem cell quiescence. Having established the diverse characteristics of healthy colonic stem cell populations, we questioned whether similar processes are present in the TME of CRC.

To address this, we performed a meta-analysis by integrating six publicly available single-cell RNA sequencing (scRNA-seq) datasets from human CRC patients, encompassing data from 82 individual donors. Following data integration (Fig 4a) and quality control (Fig 4. S1), we isolated the CSCs from the CRC samples to investigate the heterogeneity within this compartment. Clustering analysis revealed six distinct CSC populations mirroring those observed in healthy colonic tissue (Fig 4a). These clusters were initially characterised based on the similar expression of hallmark markers: CDKN1A (P21), CDKN1B (P27), CDKN1C (P57), MKI67, MUC2, and LRIG1 (Fig 4a).

**Figure 4.**
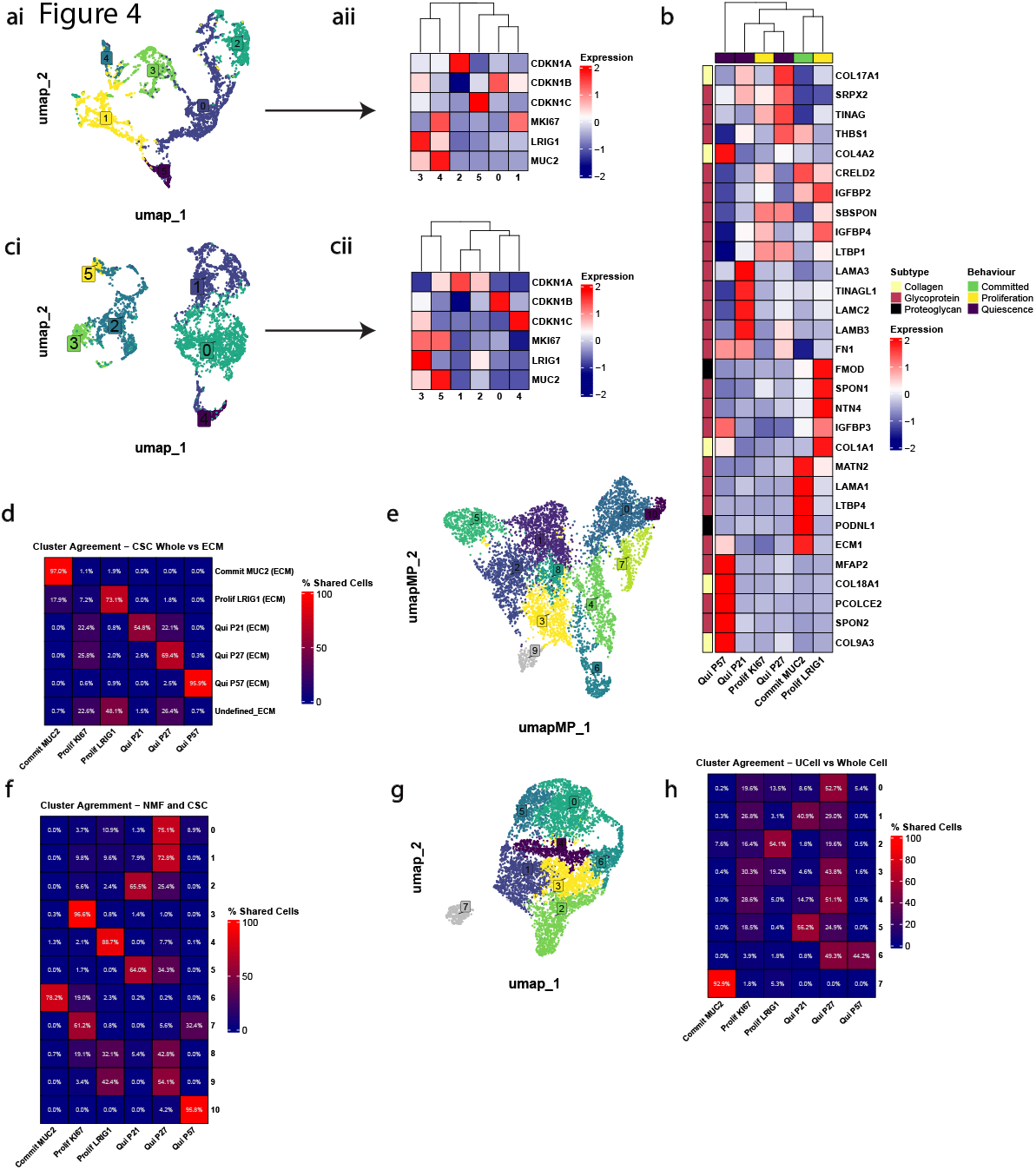
**(ai)** UMAP showing six clusters of transcriptionally unique epithelial stem cells from CRC tissue after principal component analysis (PCA) was performed. **(aii)** Clustered heatmap displaying the differentiating gene that was used for cluster annotation downstream. **(b)** Clustered heatmap of GSEA performed on annotated clusters of CSCs against the 50 Human Hallmarks gene sets. **(ci)** UMAP depicting clusters generated of CSCs from colorectal cancer when only using ECM genes as defined by the Matrisome DB. **(cii)** Clustered heatmap of the expression of the original markers that were used to annotate the clusters in **Fig 4 (ai). (d)** Agreement matrix showing the percentage of shred cells between the clusters generated using only ECM genes and the clusters generated in **Fig 4 (ai). (e)** UMAP of cells clustered using the 7 dimensional space created by the relative MP signature scores in each cell. **(f)** Agreement matrix of the percentage of shared cells between the clusters generated from the MP clustering and those in **Fig 4 (a). (g)** UMAP showing eight clusters of stem cells after using a 5-dimensional extracellular matrix signature to score against the dataset. **(h)** Agreement matrix showing the percentage of shared cells between the clusters generated using a 5-dimensional extracellular matrix signature to score and the clusters generated in **Fig 4 (a)**.

Across all six stem cell states in healthy tissues, we identified commonalities reflecting fundamental aspects of stem cell biology (Fig 4. S2a,c). 4 Notably, pathways involved in maintaining stemness and regulating the balance between quiescence and proliferation are consistently modulated. The WNT/ *β* -catenin signaling pathway, crucial for stem cell maintenance and self-renewal, is modulated according to the proliferative potential of each state, being more active in proliferative states and maintained at basal levels in quiescent states^36^. Additionally, metabolic pathways such as glycolysis and oxidative phosphorylation are adjusted based on energy demands, with lower activity in quiescent cells (Qui_P27 and Qui_P57) and higher activity in proliferative cells (Prolif_Ki67).

Again, Cell cycle regulation is modulated as would be expected, with cell cycle inhibited proteins like P21, P27 and P57 associated with down regulation of E2F Targets, G2M Checkpoint and mitotic spindle in their respective states by inhibiting cyclin-dependent kinases^37^. This downregulation corroborates their quiescent phenotype, characterised by reduced proliferation and metabolic activity, suggesting common pathway regulatory mechanisms governing theses states in both healthy and diseased tissue^38,39^(Fig 4. S2a,c).

Notably, in the Qui_P57 cluster, we observed unique enrichment of the WNT/ *β*-Catenin signaling pathway (Fig. X), which is critical for stem cell maintenance and quiescence in the intestinal epithelium^4^. The activation of WNT signaling in this cluster suggests a role in preserving the stemness and quiescent status of these cells within the TME. Additionally, pathways associated with the basement membrane, which supports stem cell niches, were enriched in the Qui_P57 cluster, indicating a potential interaction between these quiescent CSCs and their microenvironment (Fig 4. S2a,c).

In contrast, the proliferative clusters (Prolif_Ki67 and Prolif_LRIG1) displayed upregulation of pathways promoting cell cycle progression and proliferation. Specifically, Prolif_Ki67 cluster showed activation of E2F Targets, G2M Checkpoint, MYC Targets and KRAS Signaling pathways (Fig 4. S2a,c), reflecting active cell division and metabolic activity^39,40^. The expression of MKI67 further substantiates the highly proliferative nature of this cluster^41^. Notably, Prolif_LRIG1, consistent with healthy tissue, showed levels of activation through PI3K/AKT/mTOR in addition to activated KRAS, potentially oncogenic in nature. Intriguingly, this cluster shows reduced proliferative signature when compared to healthy tissue with down regulation of proliferative hallmarks like G2M E2F targets and Myc (Fig 4. S4a,c).

Using these initial findings, it is possible to see general how manner the for various quiescent pathways and are proliferative modulated CSC in populations a regardless of whether they are in a normal or diseased state. However, when the specific characteristics of these regulatory networks are compared in more detail, a more complicated picture is revealed (Fig 4. S2), where the oncogenic pathways and metabolic programs that diverge or overlap with their healthy counterparts are considered through comparative analysis.

As ECM factors have been shown to be influential on all hallmarks of cancer including cell behavior of the TME^42^, we again explored whether it was similarly linked to the cell behaviors we observed in the CSC compartment of CRC. Exploring this we conducted differential gene expression analysis, shortlisted these genes against the reference list from the MatrsiomeDB and examined them using a superimposed volcano plot grouping quiescent clusters and proliferative/committed clusters (Fig 4b, c, Fig 4. S2b). It appears that an upregulation in ECM genes *LAMA5, LAMA3, MFAP2* are show specific modulation in quiescent CSC with an inverse regulation of *MFAP2* in Proliferating/Committed CSCs. Consistently, *COL17A1*^19^ has previously been reported to be critical to regulate quiescence in LGR5+P27+ CSCs in colonic tumors and notably, we see this finding is replicated here (Fig 4b, c, Fig 4. S2b,d). In contrast to healthy tissue, *LAMA1* was more strongly associated with Muc2 cell states potentially contributing to a pro-survival signal within this oncogenic CSC subpopulation. We performed a Venn analysis comparing the shared and different up and down regulated ECM genes in each of the annotated clusters. Surprisingly between healthy and cancerous upregulated ECM genes, only the P57 clusters shared more than 20 percent of their upregulated ECM genes (Fig 4. S2e) and interestingly the shared downregulated ECM genes were less than 8 percent across all conditions. The difference in ECM genes from healthy to cancerous cells may implicate the dynamic nature of the ECM and the fact that its remodeling is implicated in many oncogenic processes. It remains to be understood whether various ECM combination can elicit common cell state activation. Considering this, it may not be surprising to observe differences in the ECM genes in CSCs compared to stem cells in the healthy counterpart.

Within the healthy colon we tissue, we identified ECM genes that successfully described various states of stem cells. Though direct cluster to cluster identity was not observed, we noted that of the 264 core ECM genes detected across healthy and cancer datasets, 66 genes and 70 genes were differentially regulated in healthy and cancer datasets respectively, of which 49 DEGs (or 5∼7% core ECM DEGs detected) were regulated in both datasets. Accordingly, we again postulated that in CRC CSCs, the same effect could be at play and ECM genes would be a reliable method of discerning the heterogeneity of the stem cell population. Accordingly, we reintegrated our data using only ECM genes and clustered our cells and defined 6 distinct sub populations of CSCs (Fig 4ci). Five of these were directly associated with the previous behavior markers we observed in the GCA dataset; *CDKN1A (P21), CDKN1B (P27), CDKN1C (P57), MUC2, LRIG1* (Fig 4cii). Interestingly, we again observed that ECM was shown to not be able to cluster the original *Prolif KI67* cluster defined by *MKI67 (KI67)*. Therefore, we annotated our new clusters based on these expressions to these clusters to be *Qui P21 (ECM), Qui P27 (ECM), Qui P57 (ECM), Commit MUC2 (ECM), Commit LRIG2 (ECM)* and a non-discriminate *ECM*. After establishing the determinants of these ECM based clusters, we wanted to test the validity of this approach in CRC based CSCs, we calculated the percentage of shared cells from our ECM only clustering method and our whole assay regular clustering method. Again, we observed excellent agreement in every subgroup of CSCs except for the proliferating/*Prolif KI67* group (Fig 4d).

Consistent with our previous methods in the healthy colonic stem cells, we used a gene discovery programme based on non-negative matrix factorisation and discovered 7 gene metaprogrammes (MP) that can be used to describe the heterogeneity we previously observed in the CSCs (Fig 4. S3a). Next, we scored each MP against our original clusters to see their relatedness to our previous results (Fig 4. S3b) and with overlap gene set enrichment analysis of hallmark biological processes (Fig 4. S3c): MP1 which can be defined by increased *E2F Targets* and *G2M checkpoint* hallmarks is enriched in the *Prolif KI67 cluster*. MP2, MP5 and MP6 are associated with are quiescent cluster *Qui P21, Qui P57* and *Qui P27*. Whereas MP3, MP4 and MP7 are most associated with the *Commit LRIG1* and *Commit MUC2* clusters defined by hallmarks inclusive of *Epithelial Mesenchymal Transition* suggesting a possible lineage to a differentiated cell type.

As previously, we further clustered our date in MP space, yielding 11 distinct clusters (Fig 4e), with distinct expression patterns of the cell state canonical markers (Fig 4. S4a). We next undertook a ranked gene set enrichment analysis and similar to healthy tissue, when comparing the MP clusters with the original clustering, consistent alignments were observed between corresponding subpopulations (Fig 4f, Fig 4. S4a,b). Qui P57 closely matched MP cluster 10, both showing reduced cell cycle activity (E2F targets, G2M checkpoint) and increased hypoxia signatures. Similarly, Qui P27 aligned with MP clusters 0 and 1, also featuring downregulation of cell cycle hallmarks and enrichment in hypoxia pathways. Again, Qui P21 showed the most activation of the quiescent population and was enriched in MP clusters 2 and 5, reflecting a transitional state with MP cluster 5 was negatively enriched for core cell cycle programs, while MP cluster 2 retained E2F target enrichment but showed reduced G2M checkpoint activity.

Once more the proliferative subpopulations, Prolif KI67 matched MP clusters 3 and 7, and were characterised by strong E2F targets, G2M checkpoint, and mitotic spindle enrichment. Conversely, Prolif LRIG aligned with MP clusters 4, 8, and 9, which lacked strong cell cycle hallmark enrichment but displayed upregulation of PI3K/AKT/MTOR signaling, glycolysis, and fatty acid metabolism, consistent with the Warburg effect previously noted in Prolif LRIG1 cells^43^.

Finally, the committed Commit MUC2 cluster closely mirrored MP cluster 6, which was enriched in E2F targets and G2M checkpoint hallmarks, confirming the original characterisation. These relationships are detailed further (Fig 4. S4c).

Having illustrated through multiple methods an ability to define CSC heterogeneity in CRC, we sought to consolidate a new specific ECM signature that could be used to define the various CSC states that would be inherently different from the healthy counterpart. Following the same methodology we had with healthy cells, we generated a Ucell ECM based signatures to describe 5 signatures (*P21*.*Sig_UCell, P27*.*Sig_UCell, P57*.*Sig_UCell, MUC2*.*Sig_UCell, LRIG1*.*Sig_UCell*). Clustering our data against the 5 dimension Ucell ECM space, and assessing the agreement whith whole transcriptome clustering demonstrated that there was excellent agreement of shared cells between ECM UCell Signature expression and our original *Qui P21, Qui P57, Commit MUC2* and *Commit LRIG1* (Fig 4g,h). While the *P27*.*Sig_Ucell* was less specific, it still had its highest expression in each of the quiescent clusters (Fig 4h, Fig 4. S5a). Encouragingly, Ucell Signature scores were discretely localised to specific clusters when evaluating the cells clusters using ECM only DEGs or those clustered in via MP signature clusters (Fig 4. S5b,c,d), demonstrating that the limited ECM signature generated can describe the various stem cell state we detected effectively.

### Cancer stem cell sub-niche regions apparent in colorectal cancer resections

Our results from scRNA-seq have suggested that gene signatures derived from CSCs in CRC may be able to predict different stem cell states and we hypothesised could result in denoting regional tissue specific areas of micro-niche environments that are enriched for a particular stem cell state over another. Spatial transcriptomics offers new methods by which we can visualise gene expression across a tissue aiding our understanding of these micro-niche environments. Accordingly, we analysed a naive visium spatial gene dataset of human colorectal cancer samples sourced from the 10X Genomics database (full details in methods) and calculated a UCell scores according to the ECM and MPs signatures we generated previously. Concentrating on stem cells, as we have previously, we selected *EPCAM*^*high*^*/LGR5*^*high*^*/CEACAM7-* regions of interest (Fig 5a, Fig 5. S1a).

**Figure 5.**
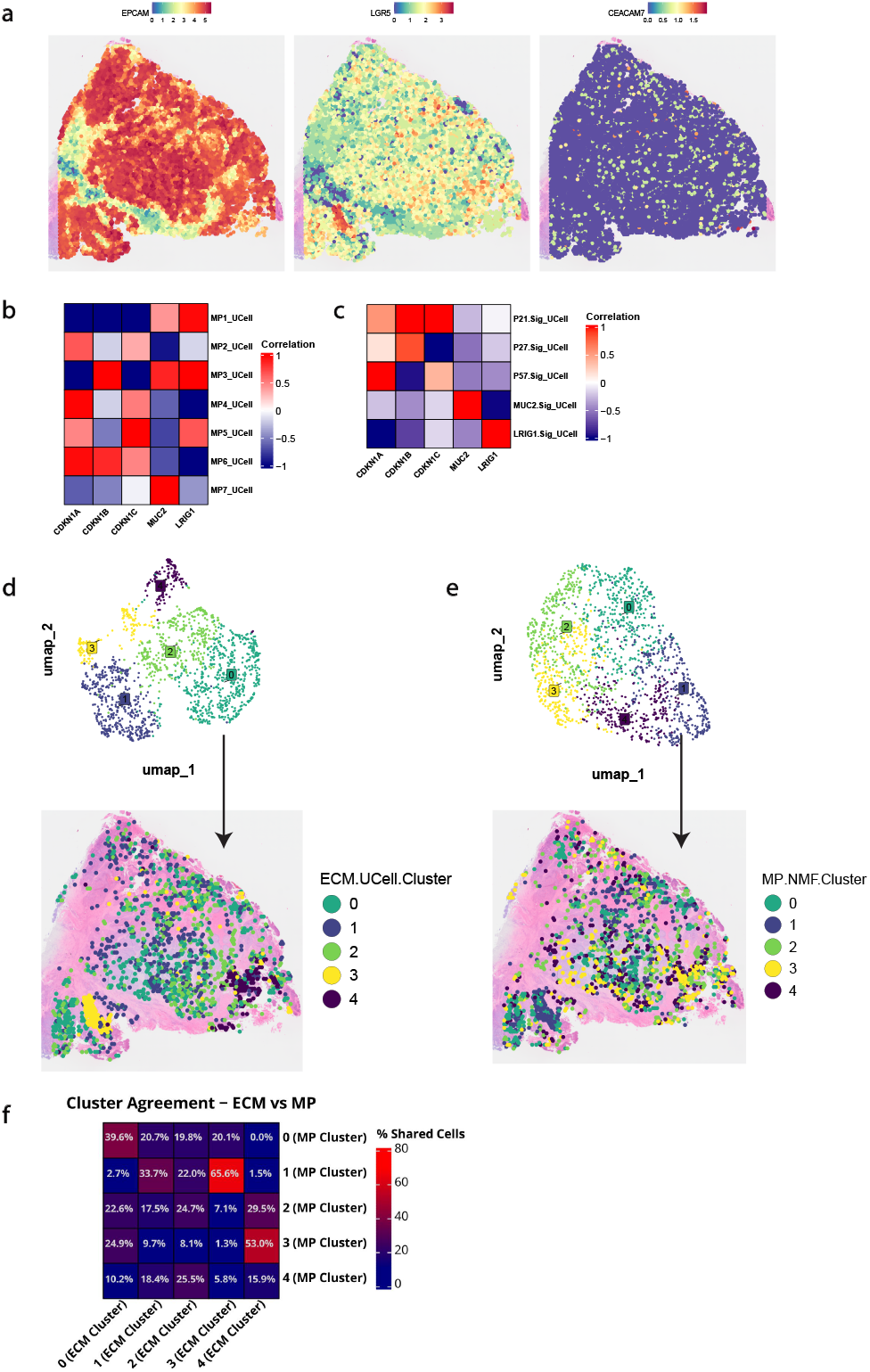
**(a)** Spatial DimPlots showing the expression of EPCAM, LGR5 and CEACAM7 across every region of interest (n = 9080) in the colorectal cancer tumour tissue section. **(b)** Heatmap that shows the correlation between the MP molecular signatures that were generated to describe various stem cell states in colorectal cancer resection, and the expression of the markers CDKN1A/P21, CDKN1B/P27, CDKN1C/P57, LRIG1 and MUC2. **(c)** Heatmap that shows the correlation between the extracellular matrix molecular signatures that were generated to describe various stem cell states in colorectal cancer resection, and the expression of the markers CDKN1A/P21, CDKN1B/P27, CDKN1C/P57, LRIG1 and MUC2. **(d)** UMAP showing five clusters of extracellular matrix signature distinct regions of interest in the CRC tissue section. SpatialDimPlot showing these clusters across the 1658 regions of interest overlaid on the CRC tissue section. **(e)** UMAP showing five clusters of MP signature molecularly distinct regions of interest in the CRC tissue section. SpatialDimPlot showing these clusters across the 1658 regions of interest overlaid on the CRC tissue section. **(f)** Agreement matrix showing the percentage of shared cells between the clusters generated using only ECM genes molecular signatures and the clusters generated using MP molecular signatures.

Clustering the signal from these ROIs using either ECM or MP signature scores revealed 5 distinct clusters in each method (Fig 5d,e). Overlaying the resultant clusters on the tissue section revealed non random distributions and concordance on the cluster grouping type across Ucell ECM or MP outputs (Fig 5f). Consistently, behavioral genes we have used throughout showed strong correlation between the ROIs’ expression of *CDKN1A*(*P21*), *CDKN1B*(*P27*), *CDKN1C*(*P57*), *MUC2* and *LRIG1* with the ECM gene signatures and MP signatures (Fig 5b,c). As previously observed, *MP2* and *MP6* displated robust G0 characteristics whereas, *MP1* and *MP7* strongly correlated with committed gene expressions (Fig 5b,c).

These results were reinforced by the visualisation of the expression in the UMAP clustering space they helped to create where the crossover in the correlation over multiple gene expressions can be seen in the regions that have the highest expression score per gene signature in both the ECM list and the MP list respectively (Fig 5. S1c). After our findings revealed that our ECM signatures effectively described quiescent gene expression of *CDKN1A* (*P21*), *CDKN1b* (*P27*) and *CDKN1C* (*P57*), and similarly these were described by *MP2* and *MP6* as a tertiary control we wanted to explore the relationship between the ECM signature expression and the MP signature expression in the regions, we wanted to further explore the relationship between the ECM gene signature and the MP gene signatures in the ROIs and therefore we calculated the correlation between them and indeed we saw that *MP2* and *MP6* were had a strong positive correlation with the quiescent signatures *P21*.*Sig, P27*.*Sig* and *P57*.*Sig* (Fig 5. S1b) supporting our earlier findings and the ability of both ECM signatures and MPs to define CSC heterogeneity.

### Pancreatic stellate cells display quiescent heterogeneity akin to colonic stem cells

Pancreatic stellate cells (PSCs) are akin to colonic epithelial stem cells, in that they oscillate between quiescent and activated states and play a crucial role in maintaining the structural integrity and function of the pancreas^44^. Given inherent similarity between the state oscillations and that these are fundamental and ancient cellular processes, we hypothesised the ECM signatures developed in healthy colon may be able to predict the various states of stellate cells. Accordingly, we analysed a publicly available CosMx™ Human Pancreas Spatial Molecular Imager (SMI) dataset that contained their CosMx™ Human Whole Transcriptome Panel. First, using the original annotation supplied with the dataset we isolated all stellate cells and performed regular downstream processes and clustered to reveal two distinct clusters of cells (Fig 6a, Fig 6. S1a). When overlaying the original annotation, we were able to see that one cluster was dominated by quiescent stellate while the other was a mixed population of both active and quiescent stellate cells (Fig 6. S1a). Next, we calculated the percentage of cells in each cluster that were quiescent or active. Encouragingly we saw an enrichment of quiescent stellate cells in annotated clusters of quiescent stellate cells (Fig 6b). We then explored these clusters for the makers of quiescence *CDKN1A, CDKN1B* and *CDKN1C* and observed their enrichment in the quiescent cluster while markers of active stellate cells were enriched the active population (Fig 6c). After establishing our own ability to differentiate these cell types we then calculated a UCell score for our five ECM gene signatures and used these data to cluster the stellate cells in 5 dimensions (Fig 6d, Fig 6. S1b,c). We investigated these clusters for specific enrichment of our ECM signatures (Fig 6e, Fig 6. S1c,d) and annotated these highly specific clusters. We also observed a positive correlation between our quiescent ECM gene signatures and the density expression of *CDKN1A, CDKN1B* and *CDKN1C* (Fig 6. S1c).

**Figure 6.**
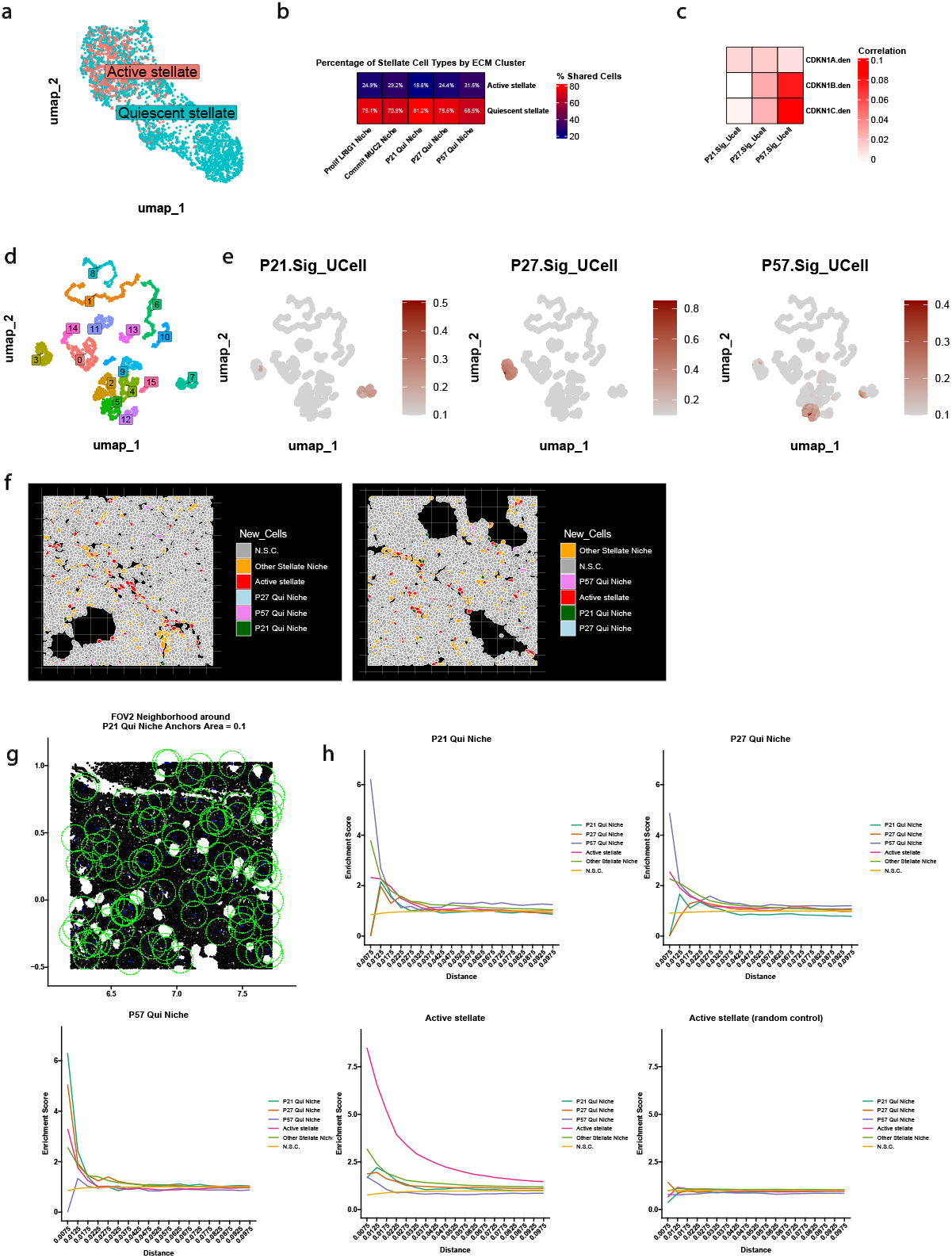
**(a)** UMAP showing two distinct clusters of pancreatic stellate cells. **(b)** Heatmap of the percentage of stellate cell type that comprise each of the five annotated groups. **(c)** Heatmap that shows the correlation between the extracellular matrix molecular signatures that were generated to describe various stem cell states in colorectal cancer resection, and the density of expression of the markers CDKN1A/P21, CDKN1B/P27 and CDKN1C/P57. **(d)** UMAP showing fifteen clusters of quiescent pancreatic stellate cells based on clustering using the healthy intestinal ECM based molecular signatures. **(e)** Feature plots of the expression of the three quiescent ECM signatures of the quiescent stellate cells. **(f)** Representative ImageDimPlots of the annotated stellate cells along with other cell types in the pancreatic tissue. **(g)** Scatter plots of the location of each cell type within the pancreas tissue field of view (FOV)2. Anchor cells are coloured in blue with the maximal radial distance shown in the green dotted circle surrounding each anchor cell. **(h)** Neighborhood analysis plots that shows the enrichment of each cell type above the background frequency surrounding each anchor cell type at increasing distances up to 0.1 mm.

Considering the robustness of the quiescent signal in this system, we reasoned we may be able to annotate quiescent domains in tissue from this system and thus, took forward only these stellate cells for further investigation (Fig 6e). Using this information, we annotated these cells to be *P21 Qui Niche, P27 Qui Niche, P57 Qui Niche* and designated all other quiescent stellate cells to be *Other Stellate Niche*. Next, taking the entire stellate cell population, we then appended the sub annotations of *P21 Qui Niche, P27 Qui Niche, P57 Qui Niche, Active stellate cells* and all *non-stellate cells (N*.*S*.*C*.) (Fig 6f, Fig 6. S1e). Subsequently, we qualitatively observed that regions of interest (ROIs) defined by CSC ECM molecular signatures were enriched within the pancreatic tissue (Fig 6f, Fig 6. S1e). The higher resolution provided by the CosMx™ SMI dataset enabled a quantitative assessment of the local microenvironment surrounding each cell.To this end, a neighborhood analysis was performed, anchoring on each of the three annotated quiescent stellate cell types, as well as the Active Stellate cells, to determine whether the pancreas contains inherent spatial micro-niches enriched for these phenotypes. Distances were measured at increasing radii up to 0.1 mm, and all counts were normalised against the background frequency of each cell type. Each anchor and the maximal radial distance across the pancreatic tissue were visualised using scatterplots (Fig 6g,h, Fig 6. S1f,g).

The immediate neighborhoods of P21 Qui Niche and P27 Qui Niche stellate cells were found to be highly enriched for P57 Qui Niche stellate cells, with enrichment levels decreasing as the distance from the anchor cells increased (Fig 6g,h, Fig 6. S1f,g). A similar pattern emerged for neighborhoods anchored on P57 Qui Niche cells, which were most enriched for P21 Qui Niche and P27 Qui Niche cells. Likewise, the Active Stellate cell population showed the greatest enrichment in neighborhoods anchored on Active Stellate cells, indicating that distinct quiescent micro-niche environments exist that may be influenced by ECM gene expression (Fig 6g,h, Fig 6. S1f,g).

To ensure these findings represented genuine biological phenomena rather than random occurrences, a control analysis was conducted. Each cell was randomly reassigned a new identity, and the neighborhood analysis was repeated. Under these randomised conditions, no enrichments above background frequency were observed (Fig 6g,h, Fig 6. S1f,g). These results confirm the presence of ECM-based molecular signature-defined quiescent stellate cell subpopulations residing in distinct, quiescent-enriched micro-niche environments within the pancreas.

## Discussion

Using a combination of in vitro culture and in vivo analysis, the present study reveals a previously underappreciated complexity in quiescent, proliferative, and lineage-committed epithelial stem cell states of the colon, thus expanding our understanding of the dynamic and heterogeneous nature of adult stem cell populations in the colon. Leveraging single-cell and spatial transcriptomic datasets, we identified and characterised at least three distinct quiescent stem cell subtypes, alongside proliferative and lineage-biased intermediates. Notably, these quiescent subtypes were robustly defined by ECM-associated gene signatures alone, underscoring the ECM as a critical regulatory element in maintaining not only a generic “resting” state but also multiple discrete, niche-dependent quiescent states.

Historically, stem cells in adult tissues have been conceptualised as a relatively uniform population transitioning between quiescence and activation, expressing varying amount of cell cycle inhibitors^2^. Recent insights have suggested that quiescence may not be a binary but rather a spectrum of states, each potentially fulfilling distinct biological functions^2,4,6^. Our findings provide direct evidence for this multiplicity: different quiescent states were delineated not only by canonical cell cycle inhibitors such as *CDKN1A (P21), CDKN1B (P27)*, and *CDKN1C (P57)*, but also by unique ECM-based transcriptional signatures. In addition, we observed state-specific enrichment of pathways traditionally associated with stemness, metabolism, and environmental sensing. For instance, deep quiescence correlated with more pronounced ECM signals, Wnt/*β* -catenin engagement, and hypoxia responses, suggesting that these cells remain metabolically poised yet stably dormant. Conversely, “primed” quiescent states appeared less metabolically constrained, exhibiting subtle shifts in oxidative phosphorylation or KRAS signaling, hinting at a readiness to re-enter the cell cycle under appropriate stimuli.

A key insight from our study is that the ECM can act as a molecular code that defines and maintains distinct stem cell subpopulations within the niche. While the ECM’s role as a non-cellular cue shaping stem cell behavior is well established^45,46^, our results move beyond the accepted paradigms. Rather than merely providing a scaffold or generic signal, we show that ECM gene expression profiles are sufficient to segregate stem cell subsets into distinct quiescent or lineage-committed states. This is especially notable in light of recent demonstrations from our group and others that laminins and other ECM components can enforce stable, reversible quiescence with unique survival mechanisms^2,13^, and that ECM-driven niches can determine cell fate in both normal and pathological conditions^5,47^.

In the context of colorectal cancer (CRC), we identified that cancer stem cells (CSCs) also occupy distinct ECM-defined states, mirroring the heterogeneity observed in healthy tissue. Notably, while the overall architectural logic of the stem cell compartment persisted in the tumor microenvironment, retaining multiple quiescent states and discrete proliferative subsets, the exact ECM signatures were altered. For example, the quiescent *P27*-positive CSC population in CRC was enriched for ECM components such as *COL17A1*, previously implicated in oncogenic quiescence and chemoresistance^19^. This suggests that while the hierarchical structure of stem cell states may be conserved, the specific ECM driven signalling is context-dependent, co-opted by tumors to maintain dormant, therapy-resistant CSC reservoirs. Such dormant CSC subsets are of significant clinical interest, as they are thought to underlie relapse after treatment and to contribute to minimal residual disease.

We also demonstrated the broader applicability of ECM-based signatures beyond the colon. Projecting the ECM signatures derived from colonic stem cells onto pancreatic stellate cells, we identified analogous quiescent subpopulations. The spatial clustering of these quiescent stellate cell subsets into discrete micro-niches further supports the notion that ECM-defined molecular states represent a fundamental principle of tissue organisation. Stellate cells, known to produce ECM components themselves and modulate pancreatic fibrosis and inflammation, are critical regulators of tissue homeostasis and disease^48^. Our ability to detect similar ECM-driven quiescent states in an entirely unrelated tissue system suggests that these principles may be broadly conserved and of general biological relevance.

Our findings collectively revise the prevailing view of stem cell heterogeneity in both the healthy and diseased states. Instead of being massive, undifferentiated pools, stem cells in the colon and pancreas are organized in a sophisticated ECM-defined sub-niche landscape with distinct molecular characteristics and possible functional importance. For therapeutic strategies, this means the ECM environment and its specific components may serve as targets to modulate stem cell states selectively, either to “wake” dormant cells to enhance their sensitivity to chemotherapy or to “lock” proliferative cells into a more quiescent, less oncogenic state. Future studies using *in vitro* or *in vivo* models will be needed to functionally validate the roles of specific ECM components within these “sleepy hollow” niches and to test whether modulating these ECM signatures can alter disease progression, improve response to therapy, or enhance tissue regeneration.

## Methods

### Data Collection

Single-cell RNA sequencing and spatial transcriptomic data was collected from various online public repositories and databases listed in the Data availability section below.

### scRNA-seq data processing and quality control

Bioinformatics analyses for scRNA-seq data were performed using the Seurat pipeline in RStudio^49^. To compare samples from various sources we consolidated all gene names to a common dictionary of Ensemble BioMart gene symbols^50^. Cells with low quality transcriptomes were excluded from analysis in a consistent manner across all datasets by excluding cells with a low UMI count (<500), extreme numbers of genes detected (<200 and >12000) and those with more than 25% of mitochondrial genes. Dimensionality reduction was completed using the top 2000 variable genes and subsequently applying principal component analysis (PCA) was performed. The principal components with the most variation was determined using the inflection point in elbow plots and were used for clustering and visualisation by Uniform Manifold Approximation and Projection (UMAP).

### Batch correction and data integration

To account for batch effects in the meta-study analysis of six CRC datasets, we employed the use of the R package STACAS^51^. This pipeline employs the use of pairwise anchors based on the mutual nearest neighbours across datasets before integrating them into one batch corrected dataset. Importantly STACAS does not rescale the data to zero mean unit variance which may contribute to masking biological differences between datasets^52^.

### Stem cell confirmation and isolation

Seurat objects were filtered to contain only cells that were *EPCAM+/LGR5+* or *EPCM+/CEACAM7+* to represent stem like and differentiated like epithelial cells. Canonical features of stemness and differentiation: *LGR5* and *CEACAM7* respectively were projected onto our UMAP space and clusters were grouped as Stem Cell or Differentiated Epithelial cell (Diff ep). Stemness was confirmed using known markers of colonic stem cells; *PROM1, SOX9, LGR5, CD44*^53,54,55^ differentiated colonic epithelial cells; *CEACAM7, CEACAM1, KRT20, GUACA2A*^56^. The crypt-axis (CA) score was computed using the following genes with the final score computed by summing the normalised expression values; *SEPP1, CEACAM7, PLAC8, CEACAM1, TSPAN1, CEACAM5, CEACAM6, IFI27, DHRS9, KRT20, RHOC, CD177, PKIB, HPGD, LYPD8*^57^. Following this we isolated EPCAM+/LGR5+/CEACAM7-cells for analysis. Next we subset our data on EPCAM+/LGR5+/CEACAM7-expression and performed standard downstream scRNA-seq analysis processing and clustering which resulted in 7 distinct transcriptional clusters. Initially when investigated for their deferentially expressed genes (DEG)s, one cluster was shown to exhibit high expression of the embryonic stem cell marker *CLDN6*. Given the GCA dataset contains several fetal samples, this was expected, though the focus on this study adult rather than embryonic stem cells, therefore these cells were removed clustering was reapplied on the remaining cells.

### Cluster annotation and Description

Behaviour markers were expressed as their mean expression per cluster scaled^58^. These were then used to annotate clusters. Pathway Inference was completed by accessing the PROGENy^59^ database through Enrichr^60^. Gene set enrichment analysis was completed using the fgsea R package^61^. Differential gene expression was performed using the Seurat FindAllMarkers function and genes lists were subsetted for ECM genes as defined by the MatrisomeDB^29^. Network analysis was performed using Cytoscape^62^.

### ECM based clustering

A new assay was created in the Seurat objects and made only of ECM gene. PCA was calculated using only highly variable genes that were in the ECM assay. Regular Seurat processes were then followed to allow for clustering based on these genes and annotation was performed using the localisation of the behavioural gene expression markers which could still be accessed through the RNA assay whilst having no impact on the formation of the clustering of the cells.

### Agreement Matrices

Agreement matrices were calculated based on the percentage of shared cells between the clusters of cells in the matrix.

### Meta-programme identification

To discover gene programmes within our data we applied non-negative matrix factorisation through the use of the MultiNMF function as part of the GeneNMF package^63^. This implements a gene discovery framework and reduces the dimesnionality of the dataset in order to detect robust meta-programmes irrespective of previous clustering methods.

### UCell Gene Signature Generation an Clustering

Deferentially expressed genes (DEGs) from each of the three analysis methods were crossed-referenced using Venn analysis. The intersect list was taken and then filtered for ECM genes as per the MatrisomeDB list^29^. In the event a gene was differentially expressed in more than one list, it remained in the list where it had the larger Average Log2 Fold Change, creating an ECM based gene signature for each stem cell phenotype. The MP gene signatures are an output from the gene discovery function using the MultiNMF function from the GeneNMF package. The AddModuleScore.UCell function is part of the UCell package^64^, and is a method for scoring gene signatures in scRNA-seq datasets. It requires a gene list as an input to calculate a gene signature score based on the Mann-Whitney U statistic. Using the gene signatures generated in the above analysis, a signature score per was calculated for each cell. These scores were then used to cluster in the number of dimensions related to number of different gene signatures; five in the ECM gene signatures lists and 8 in the MP gene signatures list.

### Spatial Transcriptomics

The CRC Visium spatial transcriptomics data were initially processed using the SCTransform function from the Seurat package^65^. Regions of interest (ROIs) were subsetted similarly to previous methods for expressing high *EPCAM+/LGR5+* and low expressing *CEACAM7* ROIs. The AddModuleScore.UCell function from the R package UCell was again implemented. These scores created five dimensions, or principal components, with which to clusters. This process was repeated for the gene signatures related to the MPs used for NMF clustering. The CosMx™ whole transciptome human pancreas dataset was normalised using the SCTransform function. Stellate cells were isolated using the annotation supplied in the metadata accompanying this dataset. These stellate cells were then clustered after performing PCA based on HVGs. Known genes of active stellate cells quiescence were visualised on UMAP. Next only “Quiescent stellate” cells were isolated. The ECM gene signatures were used to calculate a UCell score to create a new Dimensionality Reduction slot in order to cluster the cells in this multi-dimensional UMAP space.

### Neighborhood enrichment analysis

Neighborhood enrichment analysis of quiescent and active stellate cells was quantified using the nn2 function from the RANN R package. This provides fast nearest neighbour searches and calculates the abundance of cell types as a function of increasing distance from stellate cells against the background frequency of the cells^66^.

## Data availability

The single cell RNA sequencing data for the healthy colon was downloaded from the Gut Cell Atlas website. All data from this study are freely available online without restriction. The single cell RNA sequencing data for CRC can be accessed in the Gene Expression Omnibus through accession numbers GSE121455, GSE132257, GSE132465, GSE144735, GSE161277 and through Synapse under the accession code syn26720761. The CRC spatial gene expression data was accessed through the 10X Genomics Datasets by searching “colorectal cancer” and filtering software for “Space Ranger” and Species for “Human”. The human pancreas CosMx™ Spatial Molecular Imager dataset is accessible at Nanostring CosMx SMI Datasets and selecting Human Pancreas (RNA).

## Code availability

Scripts will be made available on Github upon publication.

## Notes

### Competing Interest Statement

The authors have declared no competing interest.

