## Supplemental Information for "ECM Signatures Reveal Quiescent Stem Cell Diversity in the Colonic Niche"

**77.** Moafian, Z., Maghrouni, A., Soltani, A. & Hashemy, S. I. Cross-talk between non-coding rnas and pi3k/akt/mtor pathway in colorectal cancer. *Mol. biology reports* **48**, 4797–4811 (2021).

**78.** Chen, S., Lewallen, M. & Xie, T. Adhesion in the stem cell niche: biological roles and regulation. *Development* **140**, 255–265 (2013).

**79.** Schulz-Heddergott, R. *et al.* Therapeutic ablation of gain-of-function mutant p53 in colorectal cancer inhibits stat3-mediated tumor growth and invasion. *Cancer Cell* **34**, 298–314 (2018).

**80.** Jackson, S. P. & Bartek, J. The dna-damage response in human biology and disease. *Nature* **461**, 1071–1078 (2009).

LaTeX formats citations and references automatically using the bibliography records in your .bib file, which you can edit via the project menu. Use the cite command for an inline citation.

### Acknowledgements (not compulsory)

Acknowledgements should be brief, and should not include thanks to anonymous referees and editors, or effusive comments. Grant or contribution numbers may be acknowledged.

### Author contributions statement

Must include all authors, identified by initials, for example: A.A. conceived the experiment(s), A.A. and B.A. conducted the experiment(s), C.A. and D.A. analysed the results. All authors reviewed the manuscript.

### Additional information

To include, in this order: **Accession codes** (where applicable); **Competing interests** (mandatory statement). The corresponding author is responsible for submitting a competing interests statement on behalf of all authors of the paper. This statement must be included in the submitted article file.

#### SI Note Fig 1 S2: CSC state signalling landscape

Stem cell behaviors, encompassing self-renewal, differentiation, quiescence, and survival, are governed by intricate signaling networks derived from both intrinsic and extrinsic sources. Canonical pathways such as Wnt, Notch, TGF-$\beta$/BMP, Hedgehog, JAK/STAT, mTOR, p53, and signals reducing apoptotic potential have been widely implicated in maintaining stem cell niches and regulating stem cell fate across various tissues[4,67–71]. Historically, these pathways have been considered as acting uniformly within an adult stem cell population. However, accumulating evidence suggests that stem cells do not exist as a single, homogeneous pool. Instead, they occupy a spectrum of states, from deeply quiescent to actively proliferating and on to lineage-committed cells, that may engage different aspects of these signaling pathways.

**Quiescent Stem Cells (Qui P21, Qui P27, Qui P57)** Quiescent stem cells are often defined by low levels of cell cycle progression and elevated expression of CDK inhibitors such as p21, p27, and p57[2]. These features limit metabolic activity and cell cycle entry while promoting long-term survival and genomic stability. Our analysis highlights three distinct quiescent subsets with differing degrees of dormancy. All three quiescent subpopulations (Qui P21, Qui P27, Qui P57) show reduced glycolysis, while Qui P21 and Qui P57 additionally downregulate oxidative phosphorylation. Qui P57 stands out as the most deeply dormant, displaying broad suppression of both proliferative and metabolic pathways, as well as major protein synthesis signals like PI3K/AKT/mTOR. Moreover, Qui P57 exhibits the highest Wnt activity, signalling associated with stemness maintenance, indicating that this subset maintains a stable, deeply quiescent state while remaining poised for future activation.

Qui P27 appears moderately quiescent, retaining some oxidative phosphorylation and showing pro-activation KRAS signaling, suggesting a more metabolically prepared state than Qui P57. Qui P21 represents a “primed” quiescence, downregulating cell cycle progression while simultaneously upregulating DNA repair, KRAS, PI3K/AKT/mTOR, and ROS pathways. This profile suggests Qui P21 cells are on the cusp of re-entering the cell cycle upon receiving appropriate stimuli. Together, the quiescent subsets present a hierarchy of dormancy, from deeply silent (Qui P57) to metabolically responsive (Qui P27) and primed for reactivation (Qui P21).

**Proliferative Stem Cells (Prolif KI67, Prolif LRIG1)** In contrast, proliferative stem cells exhibit elevated cell cycle activity and enhanced metabolic demands. Our analysis identified two proliferative subsets, Prolif KI67 and Prolif LRIG1, differing in intensity and profile of their proliferation signals. Prolif KI67 cells show robust upregulation of cell cycle-related genes (E2F targets, G2M checkpoint, MYC targets) and heightened metabolic activity (glycolysis, oxidative phosphorylation), marking a state of rapid, active proliferation. Prolif LRIG1 cells also demonstrate a proliferative phenotype, but with comparatively lower E2F, MYC, and KRAS signaling, possibly indicating a slower cycling population. These data align with emerging evidence that

**77.** Moafian, Z., Maghrouni, A., Soltani, A. & Hashemy, S. I. Cross-talk between non-coding rnas and pi3k/akt/mtor pathway in colorectal cancer. *Mol. biology reports* **48**, 4797–4811 (2021).

**78.** Chen, S., Lewallen, M. & Xie, T. Adhesion in the stem cell niche: biological roles and regulation. *Development* **140**, 255–265 (2013).

**79.** Schulz-Heddergott, R. *et al.* Therapeutic ablation of gain-of-function mutant p53 in colorectal cancer inhibits stat3-mediated tumor growth and invasion. *Cancer Cell* **34**, 298–314 (2018).

**80.** Jackson, S. P. & Bartek, J. The dna-damage response in human biology and disease. *Nature* **461**, 1071–1078 (2009).

LaTeX formats citations and references automatically using the bibliography records in your .bib file, which you can edit via the project menu. Use the cite command for an inline citation.

### Acknowledgements (not compulsory)

Acknowledgements should be brief, and should not include thanks to anonymous referees and editors, or effusive comments. Grant or contribution numbers may be acknowledged.

### Author contributions statement

Must include all authors, identified by initials, for example: A.A. conceived the experiment(s), A.A. and B.A. conducted the experiment(s), C.A. and D.A. analysed the results. All authors reviewed the manuscript.

### Additional information

To include, in this order: **Accession codes** (where applicable); **Competing interests** (mandatory statement). The corresponding author is responsible for submitting a competing interests statement on behalf of all authors of the paper. This statement must be included in the submitted article file.

#### SI Note Fig 1 S2: CSC state signalling landscape

Stem cell behaviors, encompassing self-renewal, differentiation, quiescence, and survival, are governed by intricate signaling networks derived from both intrinsic and extrinsic sources. Canonical pathways such as Wnt, Notch, TGF-$\beta$/BMP, Hedgehog, JAK/STAT, mTOR, p53, and signals reducing apoptotic potential have been widely implicated in maintaining stem cell niches and regulating stem cell fate across various tissues[4,67–71]. Historically, these pathways have been considered as acting uniformly within an adult stem cell population. However, accumulating evidence suggests that stem cells do not exist as a single, homogeneous pool. Instead, they occupy a spectrum of states, from deeply quiescent to actively proliferating and on to lineage-committed cells, that may engage different aspects of these signaling pathways.

**Quiescent Stem Cells (Qui P21, Qui P27, Qui P57)** Quiescent stem cells are often defined by low levels of cell cycle progression and elevated expression of CDK inhibitors such as p21, p27, and p57[2]. These features limit metabolic activity and cell cycle entry while promoting long-term survival and genomic stability. Our analysis highlights three distinct quiescent subsets with differing degrees of dormancy. All three quiescent subpopulations (Qui P21, Qui P27, Qui P57) show reduced glycolysis, while Qui P21 and Qui P57 additionally downregulate oxidative phosphorylation. Qui P57 stands out as the most deeply dormant, displaying broad suppression of both proliferative and metabolic pathways, as well as major protein synthesis signals like PI3K/AKT/mTOR. Moreover, Qui P57 exhibits the highest Wnt activity, signalling associated with stemness maintenance, indicating that this subset maintains a stable, deeply quiescent state while remaining poised for future activation.

Qui P27 appears moderately quiescent, retaining some oxidative phosphorylation and showing pro-activation KRAS signaling, suggesting a more metabolically prepared state than Qui P57. Qui P21 represents a "primed" quiescence, downregulating cell cycle progression while simultaneously upregulating DNA repair, KRAS, PI3K/AKT/mTOR, and ROS pathways. This profile suggests Qui P21 cells are on the cusp of re-entering the cell cycle upon receiving appropriate stimuli. Together, the quiescent subsets present a hierarchy of dormancy, from deeply silent (Qui P57) to metabolically responsive (Qui P27) and primed for reactivation (Qui P21).

**Proliferative Stem Cells (Prolif KI67, Prolif LRIG1)** In contrast, proliferative stem cells exhibit elevated cell cycle activity and enhanced metabolic demands. Our analysis identified two proliferative subsets, Prolif KI67 and Prolif LRIG1, differing in intensity and profile of their proliferation signals. Prolif KI67 cells show robust upregulation of cell cycle-related genes (E2F targets, G2M checkpoint, MYC targets) and heightened metabolic activity (glycolysis, oxidative phosphorylation), marking a state of rapid, active proliferation. Prolif LRIG1 cells also demonstrate a proliferative phenotype, but with comparatively lower E2F, MYC, and KRAS signaling, possibly indicating a slower cycling population. These data align with emerging evidence that

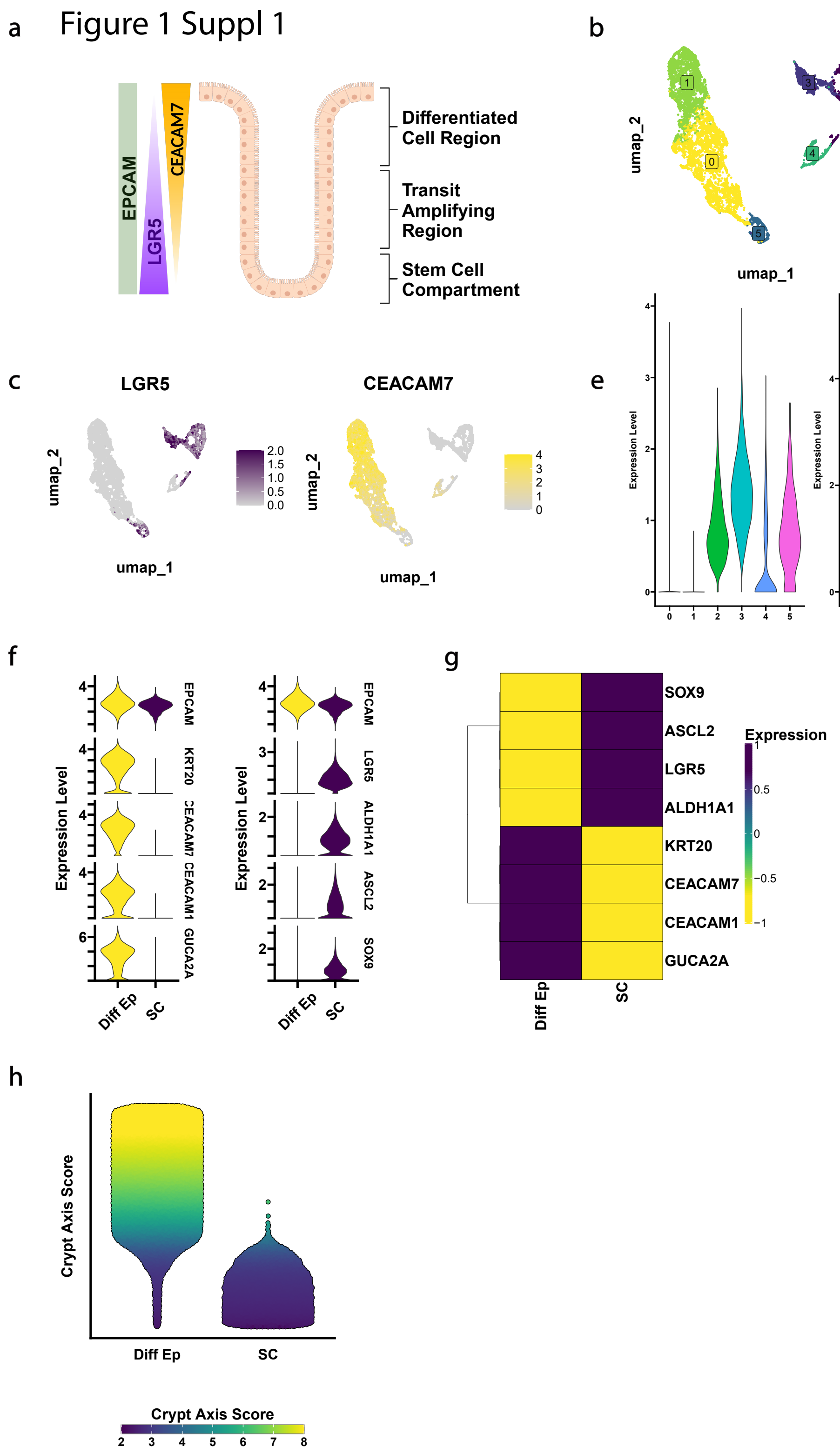


**Figure 1. S1.** **(a)** Overview of the colonic crypt-axis separated by biological zone and the expected expression gradients for canonical epithelial marker EPCAM, canonical stem marker LGR5, and canonical differentiated epithelial cell marker CEACAM7. **(b)** UMAP of six clusters identified in a population of EPCAM+/LGR5+/CEACAM7+ cells. **(c)** Feature plots representing the cellular expression of LGR5 in the UMAP clustered space. **(d)** Feature plots representing the cellular expression of CEACAM7 in the UMAP clustered space. **(e)** Violin plots showing the expression of genes LGR5 and CEACAM7 in each cluster. **(f)** Violin plots comparing various known genes associated with epithelial cell differentiation and stemness between differentiated epithelial (Diff Ep) and stem cells (SC). **(g)** Clustered heatmap of scaled expression between Diff Ep and SC. **(h)** Bee swarm plot of the crypt-axis score between the two groups where an increased score represents an increased likelihood a cell's location would be expected near the top of the crypt-axis.

LRIG1, originally proposed as a quiescence marker, actually delineates a distinct proliferative subset. This finding challenges prior assumptions and suggests that proliferative capacity can be graded within the stem cell pool.

**Committed Stem Cells (Commit MUC2)** As cells move along the trajectory toward differentiation, their signaling environment changes dramatically. The Commit MUC2 subset represents cells transitioning to a goblet cell lineage, as indicated by strong MUC2 expression. These cells show downregulation of proliferative pathways and a shift toward specialised functions, including upregulation of apical surface components and the unfolded protein response (UPR) to support high secretory demands. The overall signaling signature of Commit MUC2 cells reveals that differentiation is accompanied by a down-tuning of cell cycle activity and a shift toward pathways that facilitate specialised epithelial function.

**Reconsidering Stem Cell Signaling as a Spectrum** Our detailed analysis of quiescent, proliferative, and committed stem cell states including their respective subpopulations, highlights a more nuanced signaling landscape than previously appreciated. Rather than viewing adult stem cells through a single, uniform signaling paradigm, these data support a dynamic, hierarchical model. Quiescent states are not simply "off"; instead, they exist in varying degrees of metabolic and proliferative readiness. Similarly, proliferative states differ in their intensity of cell cycle engagement, and cells in a committed state show unique pathway activation tailored to their specialised functions.

This complexity also challenges standard interpretations of niche signaling. While widely held models suggest uniform gradients of signals like Wnt and BMP exist in the niche[4] but that theses are largely a uniform within the stem cell populations, our findings indicate that these gradients may be filtered differently by each subpopulation. Additionally, ECM composition and signalling, for example, laminin-mediated MAPK activity[13], further refine our understanding of how local niches create distinct states within the broader stem cell compartment.

Taken together, these data reveal a signaling mosaic that underpins the functional heterogeneity of colonic stem cells. By moving beyond a one-size-fits-all approach, we can now appreciate a dynamic compartment where deep dormancy, varying levels of quiescence, active proliferation, and lineage commitment each correspond to distinct signaling environments. This refined understanding holds significant implications for tissue homeostasis, repair, and pathology, and sets the stage for more targeted interventions aimed at manipulating specific stem cell states.

### SI Note Fig 2 S1: MP Characterisation of the stem cell niche

The following supplementary material contains extended discussion on the computational results that enhance the results given in the main body of the text. As stated, the application of MPs in defining intestinal stem cell (ISC) heterogeneity showed that MP-based clustering was largely similar to initial transcriptome-based clustering approaches. These comparisons, based on cell clustering and pathway analysis, confirm the effectiveness and reliability of the MP approach in identifying the various states of stem cells in the colonic epithelium.

For example, the *Prolif KI67* cluster exhibited strong specificity to MP clusters 6 and 7, which were enriched in classic cell cycle hallmarks such as E2F Targets and the G2M Checkpoint (Fig 2. S1c). In contrast, the *Prolif LRIG1* cluster aligned most closely with MP cluster 3, suggesting a slower yet still active cell cycle, as indicated by G2M Checkpoint enrichment (Fig 2. S1c). Similarly, MP cluster 2 most closely resembled the original *Commit MUC2* cluster, again reflected by enrichment in Mitotic Spindle, G2M Checkpoint, and EMT hallmarks (Fig 2. S1c).

Conversely, the quiescent *Qui P21* cluster was most similar to MP clusters 0 and 8, both of which displayed downregulation of E2F Targets and G2M Checkpoint, coupled with enrichment in Hypoxia (Fig 2. S1c). A similar pattern emerged for the *Qui P27* cluster, which resembled MP clusters 4 and 5. Both of these clusters were negatively enriched for classic cell cycle progression hallmarks, including E2F Targets and G2M Checkpoint (Fig 2. S1c). Likewise, the *Qui P57* cluster was more similar to MP cluster 1, which also exhibited downregulation of E2F Targets and G2M Checkpoint and upregulation of Hypoxia (Fig 2. S1c).

### SI Note Fig 4 S1: Stem cell isolation in colorectal cancer epithelial cells

To isolate stem cells we again followed a similar protocol as we had in healthy tissue. As above selecting cells positive for *EPCAM+/LGR5+* or *EPCAM+/CEACAM7+* to reflect the diverse nature of epithelial cells involved in CRC from stem to differentiated. We took these cells and investigated whether the expressions of *LGR5* and *CEACAM7* were distinct and saw that indeed, expression of *LGR5* was absent in cells expressing *CEACAM7* (Fig ref). This was used to annotate the cells into two groups, CSCs and differentiated epithelial (Diff EP). We inspected our groups for their expression of known markers of stemness and differentiation inclusive of the original markers used to isolate these cells from their original datasets (*EPCAM/LGR5/CEACAM7*). Indeed, it was seen that stem cells have increased expression in stem markers *PROM1*, *CD44* and *SOX9* while Diff Ep cells have increased expression of KRT20, CEACAM1 and GUACA2A (Fig. reference). This was further investigated by determining the crypt-axis score of every cell in the CSC and Diff Ep groups. This analysis revealed a skew towards the crypt base in CSCs compared to differentiated epithelial cells (Fig. reference), suggesting our successful method of CSC isolation from CRC scRNA-seq data.

### SI Note Fig 4 S2: Comparative signalling landscape between healthy can CRC stem cell niche

Comparative analysis between healthy and cancerous tissues revealed both commonalities and differences in pathway activation among the stem cell states. In healthy tissue, the quiescent clusters maintain downregulation of proliferative pathways and upregulation of stem cell maintenance pathways, preserving tissue homeostasis. However, in CRC samples, we observed aberrant activation of oncogenic pathways within these clusters. For example, the quiescent clusters in CRC displayed upregulation of DNA Repair and IL6/JAK/STAT3 Signaling pathways (Fig. Fig 4 S2a and c), which may contribute to therapy resistance and tumor survival[72,73]. The Qui_P21 cluster had upregulation of IL6/JAK/STAT3 Signaling hallmark, as well as an increase of DNA repair, and MYC targets that was not present in the healthy counterpart. These are implicated in pro-oncogenic functions involving immune evasion and a response to genomic instability, aiding in the survival of cells with aberrant cancerous DNA mutations, and furthermore are even more primed to exit quiescence and reenter the cell cycle[40]. The Qui_P27 cluster in CRC also showed increased activation of WNT/$\beta$-Catenin, Notch, and Hedgehog signaling pathways, indicating a shift toward a more activated state and potential for tumor initiation[36,67,69]. There was also a marked reduction in this cluster of Oxidative Phosphorylation, suggesting the shift of these towards glycolysis to meet the energy demands of the hypoxic tumour environment[74]. The Qui_P57 cluster also demonstrated notable oncogenic changes compared to the healthy intestinal stem cells whereby there was a marked decrease in Interferon Alpha and Gamma Responses, indicating impaired immune surveillance[75]. This coincided with a noticeable enhancement of IL2 STAT5 Signalling hallmark which encourages cellular survival and immune evasion in cancer cells[76].

Furthermore, in the more active associated CSCs, the Commit_Muc2 cluster, associated with differentiated goblet cells, exhibited activation of oncogenic pathways such as PI3K/AKT/mTOR Signaling in CRC samples, suggesting that differentiated cells may acquire proliferative and tumorigenic properties[77]. Additionally, there was a decrease in the Apical Junction Hallmark, this suggests a reduction in the expression of cell adhesion molecules which may be disrupting the normal tissue architecture of the colon, thereby facilitating tumour growth[78]. Moreover, the Prolif_KI67 cluster had an increase in P53 pathway and Mitotic Spindle hallmarks which were not previously observed. These could reflect the increased cell division, and a response to aberrant genomic instability known to be present in CRC. Mutant P53 is known to promotes tumour growth as it cannot trigger apoptosis, enabling the uncontrolled proliferation of these CSCs[79]. Likewise, in the Prolif_LRIG1 cluster, there was a notable downregulation of DNA repair mechanisms, suggesting an increased likelihood of genomic mutation accumulation while these cells are dividing[80]. This was paired with an upregulation in Glycolysis and Fatty Acid and Cholesterol Metabolism, that was previously not observed in the healthy counterparts. Another indication of the Warburg effect in the CSC population, while the increases in Fatty Acid and Cholesterol in metabolism could be a critical component in lipid synthesis that is needed for membrane production during persisitent cellular division[43].

These findings highlight the disruption of normal regulatory mechanisms in CRC, leading to increased proliferation and survival across various stem cell states. Despite these oncogenic alterations, there appears broad concordance across healthy and diseased tissue relating to the states that stem cells may occupy, an important consideration when therapeutic efficacy is considered. Understanding these differences provides valuable insights into the mechanisms underlying CSC behavior in CRC and may inform the development of targeted therapies to overcome resistance and prevent recurrence.

### SI Note Fig 4 S4 Generation of a ECM Ucell signaute for CSC cells

As we undertook in healthy tissue, we took the intersect of the DEGs from the three methods of clustering for similar clusters (Fig ref) and subsetted out only ECM genes (FIG REF). This enabled the creation of 5 new distinct ECM signatures (*P21.Sig_UCell, P27.Sig_UCell, P57.Sig_UCell, MUC2.Sig_UCell, LRIG1.Sig_UCell*) that could be used to classify the diverse nature of the CSC population. As shown previously, we were unable to define the *Prolif KI67* cluster using ECM clustering, therefore we did not define an ECM based signature to define this phenotype. Subsequently, we calculated a UCell score for each cell based on the expression of these signatures and used these values to cluster the CSCs in a 5-dimensional space and observed 8 clusters of sub-populations of CSCs as defined by our ECM signatures (Fig ref). Independently of clustering, we wanted to examine the relationship between the expression of our ECM signatures and the expression markers we had used to annotate different CSC states. We saw strong positive correlations between *CDKN1A*(*P21*), *CDKN1C*(*P57*), *MUC2* and *LRIG1* with their respective signatures while *CDKN1B*(*P27*) was less specific but still a weak positive correlation was seen (Fig ref).

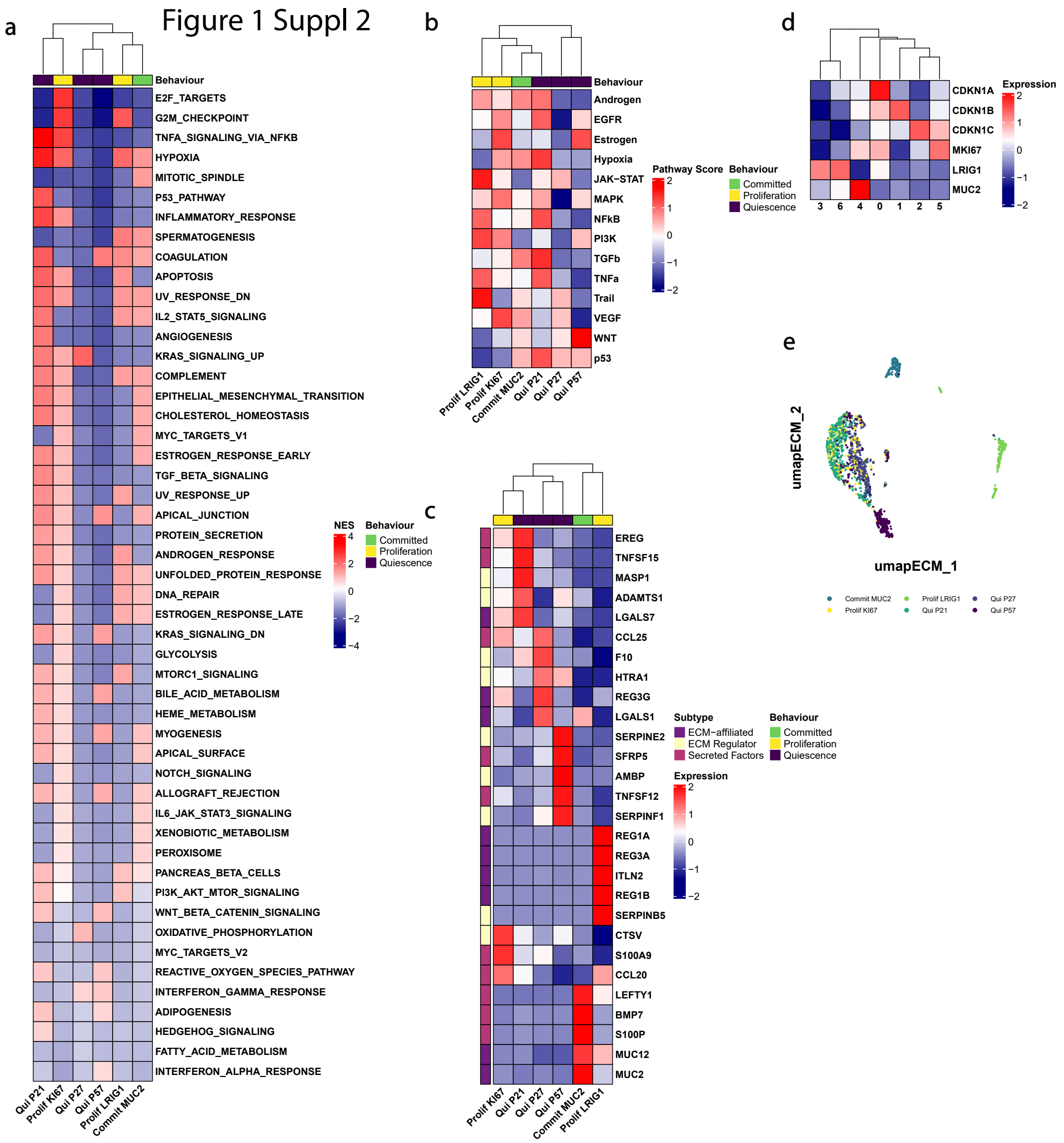


**Figure 1. S2.** **(a)** Clustered heatmap of Gene Set Enrichment Analysis performed on annotated clusters of healthy stem cell from the Gut Cell Atlas (GCA) database ran against the 50 Human Hallmarks gene sets.**(b)** Clustered heatmap of Pathway Inference Analysis performed on annotated clusters of stem cells ran the PROGENy database gene sets.**(c)** Clustered heatmap of the top 5 Matrisome Associated ECM DEGs expressed per cluster. **(d)** Clustered heatmap of the expression of the original markers that were used to annotate the clusters in **Fig. 1g**. **(e)** Visualisation of the original cells' cluster identity from **Fig. 1b** in the ECM cluster UMAP space of **Fig. 1g**.

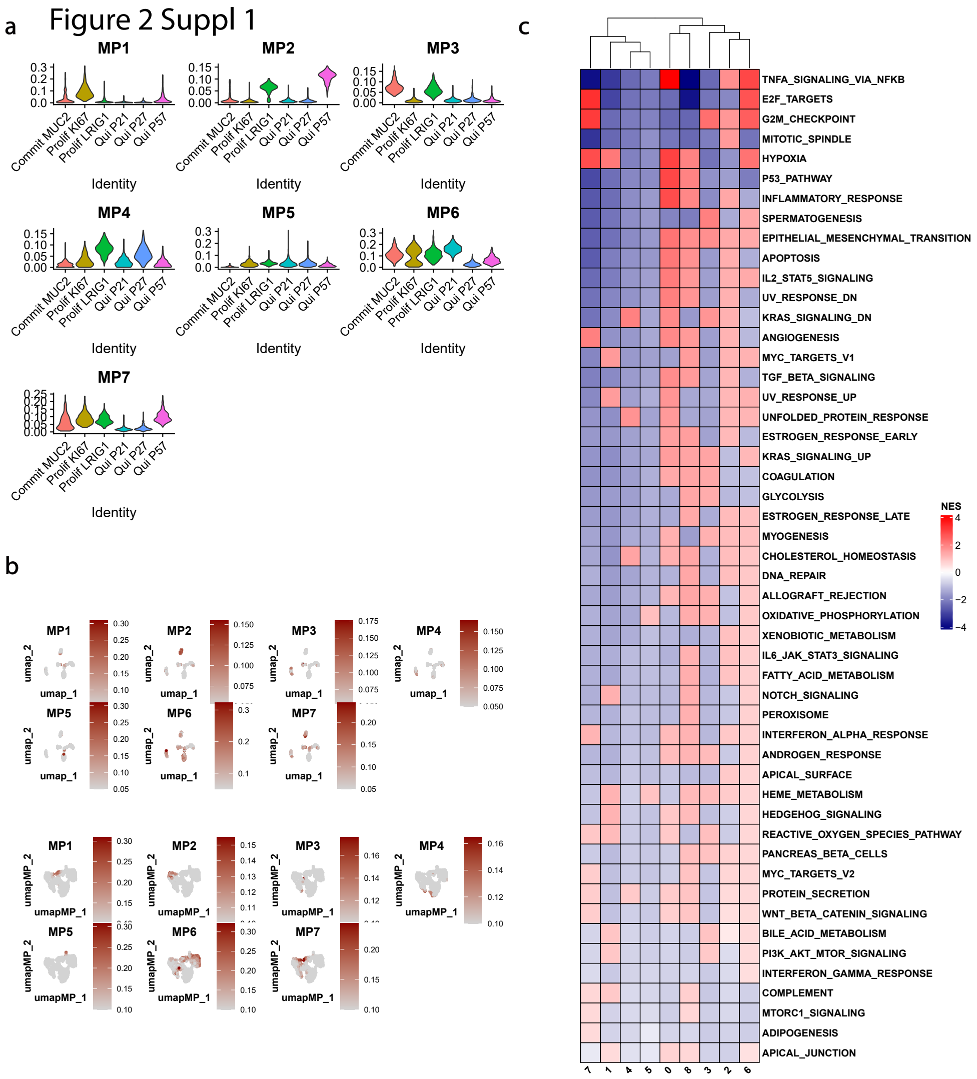


**Figure 2. S1.** **(a)** Violin plots displaying the UCell signature score in each cluster as defined in **Fig 2 (a)**. **(b)** Featureplots showing the expression of the MP signatures in the whole cell clustering and ECM clustering approaches from **Fig 2**. **(c)** Clustered heatmap of the GSEA performed against the nine clusters generated using MPs in **Fig 2**

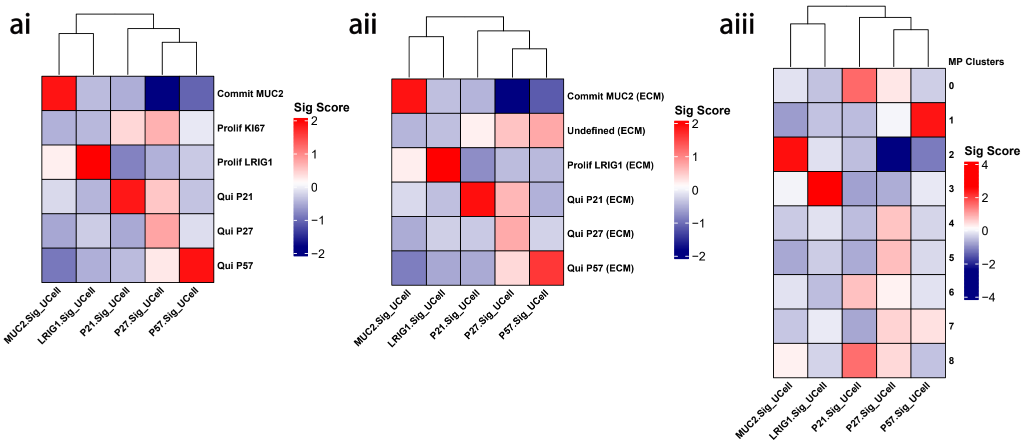


**Figure 3. S1.** **(ai)** Heatmap displaying the various ECM molecular signatures against the clusters that were generated using the whole transcriptome. **(aii)** Heatmap displaying the various ECM molecular signatures against the clusters that were generated using ECM genes only. **(aiii)** Heatmap displaying the various ECM molecular signatures against the clusters that were generated using MPs.

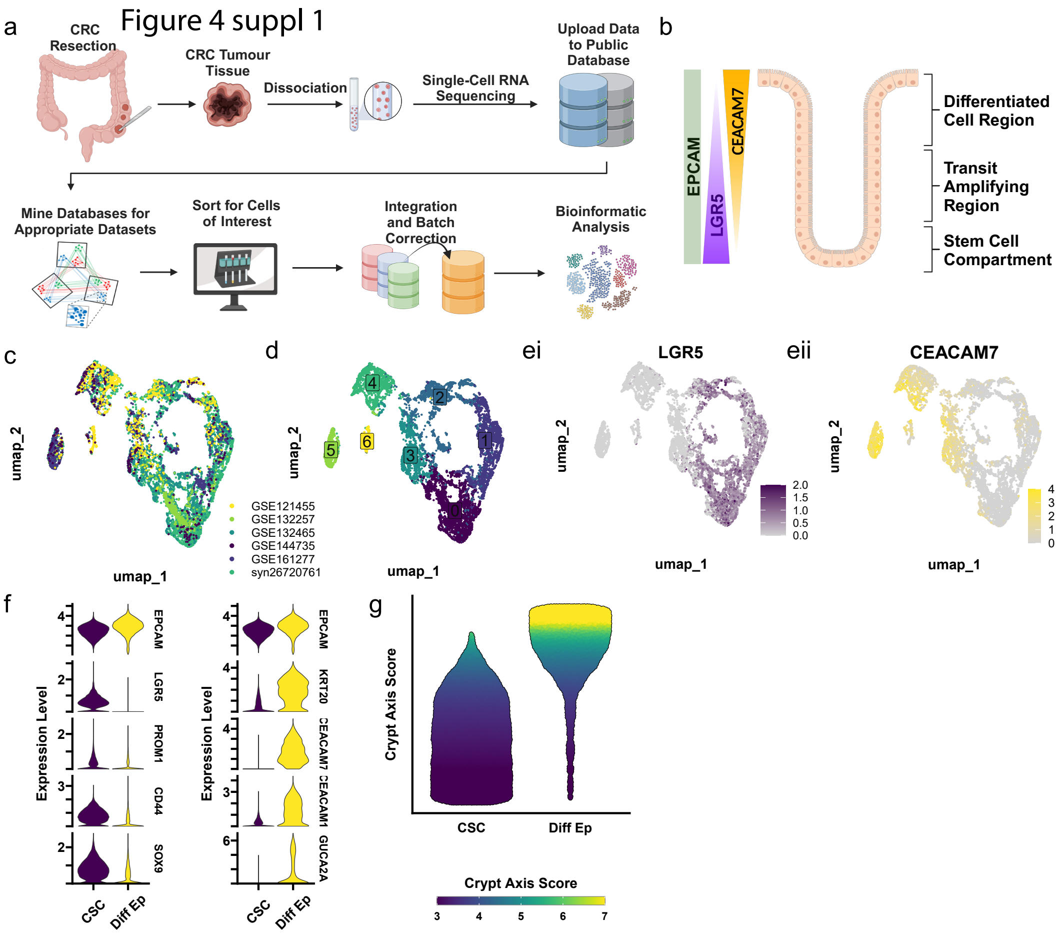


**Figure 4. S1.** **(a)** Overview of the workflow. In short: utilising publicly available single-cell RNA sequencing on human resected colorectal cancer tumours, data was collected, downloaded and integrated to correct batch effects. This allowed for appropriate bioinfomatic analyses in RStudio. **(b)** Overview of the colonic crypt-axis separated by biological zone and the expected expression gradients for canonical epithelial marker EPCAM, canonical stem marker LGR5, and canonical differentiated epithelial cell marker CEACAM7. **(c)** UMAP of six clusters identified in a population of EPCAM+/LGR5+/CEACAM7+ cells from colorectal cancer singel cell RNA sequencing data. **(d)** UMAP showing the dataset origin of each cell. Each colour relates the unique accession codes that can be used to access these datasets. **(e)** Feature plots representing the cellualr expression of LGR5 and CEACAM7 in the UMAP clustered space. **(f)** Violin plots comparing various known genes associated with epithelial cell differentiation and stemness between *differentiated epithelial* (*Diff Ep*) and *cancer stem cells* (*CSC*). **(g)** Bee swarm plot of the crypt-axis score between the two groups where an increased score represents an increased likelihood a cell being a differentiated epithelial cell expected to be near the top of the crypt-axis and a lower score would represent an increased likelihood of a cell being stem-like.

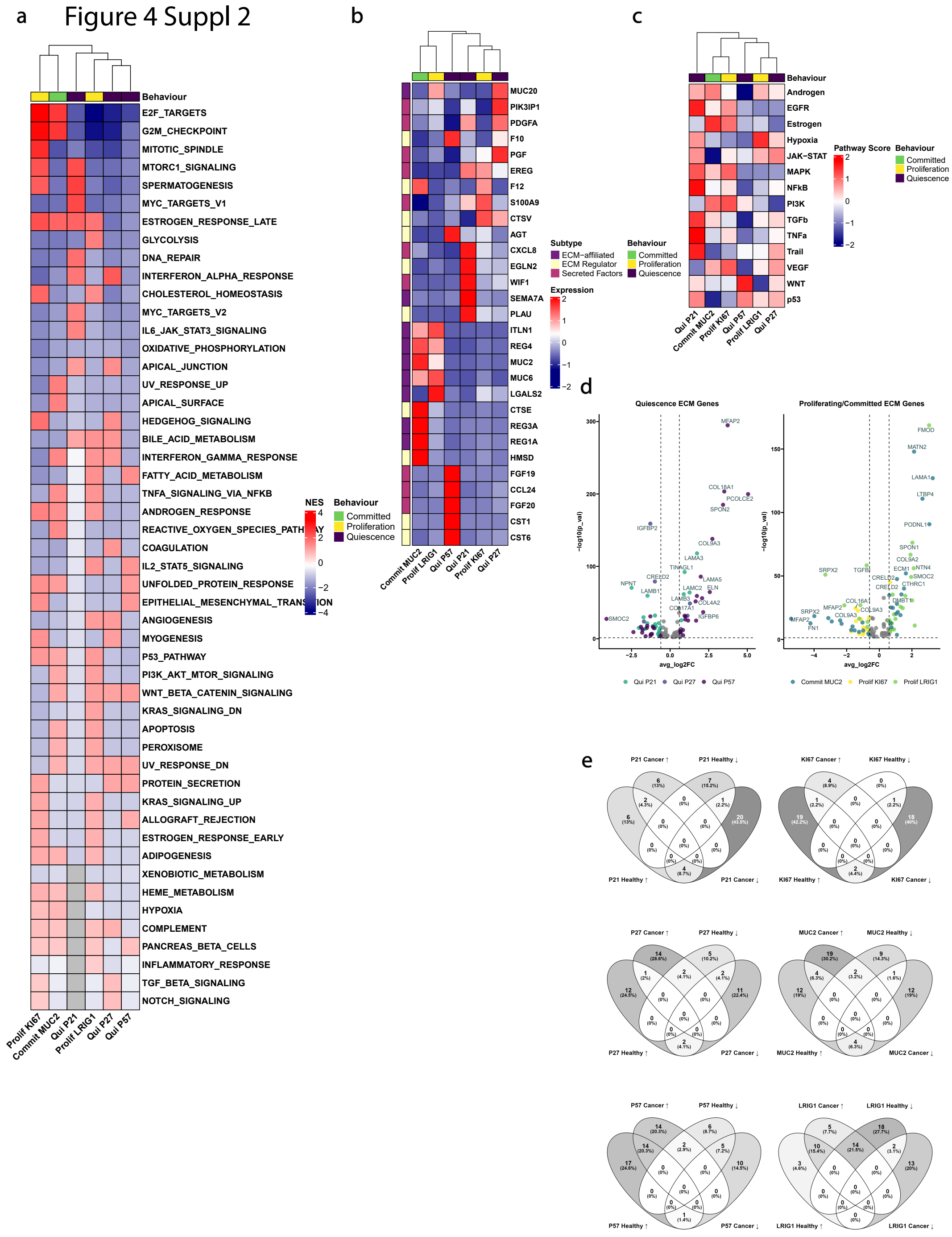


**Figure 4. S2.** **(a)** Clustered heatmap of Gene Set Enrichment Analysis performed on annotated clusters ofCSCs ran against the 50 Human Hallmarks gene sets. **(b)** Clustered heatmap of the top 5 Matrisome Associated ECM DEGs expressed per cluster. **(c)** Clustered heatmap of Pathway Inference Analysis performed on annotated clusters of stem cells ran the PROGENy database gene sets. **(d)** Volcano plot of ECM DEGs in the active-like clusters, as per their Average Log 2 Fold Change. **(e)** Venn analysis of the ECM genes only that are associated with each of the three clustering methods for the CSC dataset.

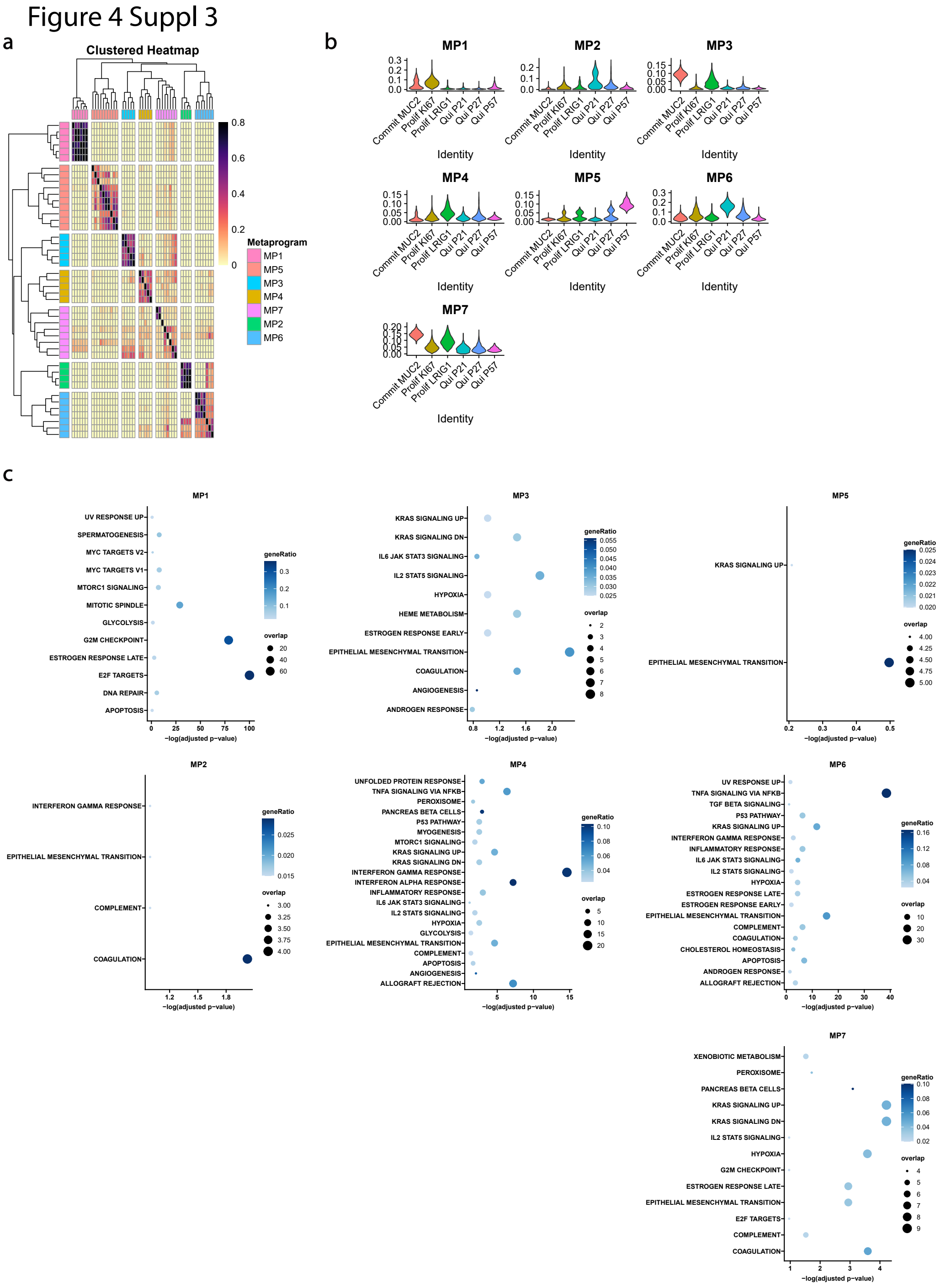


**Figure 4. S3.** **(a)** Jaccard plot of the individual gene programmes that comprise the MPs. **(b)** Violin plots displaying the UCell signature score in each cluster as defined in **Fig 4**. **(c)** Dotplot describing the GSEA Hallmarks that are described by each of the seven Meta-programmes.

Figure 4 Suppl 4

a

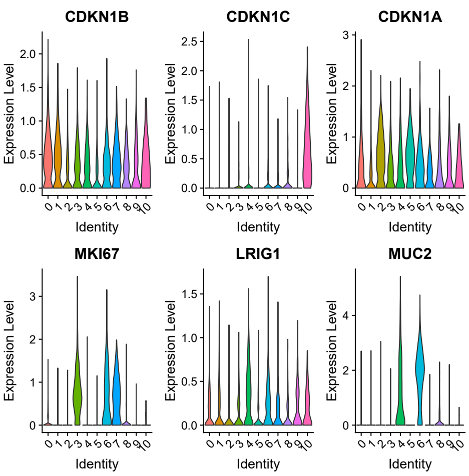


b

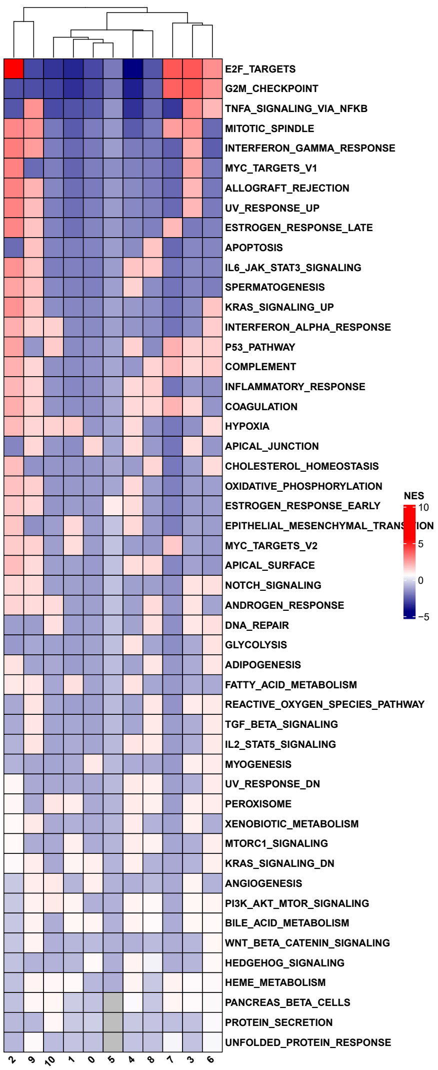


c

| Meta-Programme | Original Clusters | MP Clusters | Gene List Subset |
|---|---|---|---|
| MP1 | Prolif KI 67 | 3, 7 | ANLN, ANP32E, ARHGAP11A, ASF1B, ASPM, ATAD2 |
| MP2 | Qui P21, Qui P27 | 5 | ACSL6, ADIRF, APOA2, ARHGAP6, ATP5F1EP2, BEX2 |
| MP3 | Commit MUC2, Prolif LRIG1 | 4, 6 | AFAP1-AS1, AGR3, AHCYL2, ALDH1L1, ANO1, ATP2A3 |
| MP4 | Prolif LRIG1 | 4, 9 | ACTRT3, BAG3, BARX1, BNIP3, CASP1, CCL2 |
| MP5 | Qui P57 | 0, 4, 7, 10 | ACTL8, ATP6V0C, BEX4, C6orf15, CALD1, CALML3 |
| MP6 | Qui P21, Qui P27, Prolif KI67 | 2, 3, 5, 8, 9 | ARL14, BIRC3, CLCF1, CXCL8, DUSP5, EDN1 |
| MP7 | Commit MUC2, Prolif LRIG1 | 4, 6, 8, 9 | CPS1, LGALS2, SULT1E1, TTR, ANPEP, DEFA5 |

**Figure 4. S4.** (**a**) Violin plots of the expression of each MP score in each of the clusters generated using the MP clustering approach. (**b**)Clustered heatmap of Gene Set Enrichment Analysis performed on MP clusters of stem cells ran against the 50 Human Hallmarks gene sets. (**c**) Summary table of the relationships between the meta-programmes, the original whole cell clustering identities, the new meta-programme derived clusters and a subset of the genes that drove the original meta-programmes in **Fig 4**.

Figure 4 Suppl 5

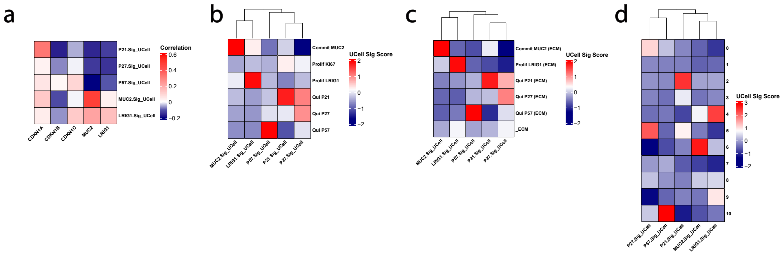


**Figure 4. S5.** **(a)** Heatmap that shows the correlation between the extracellular matrix molecular signatures that were generated to describe various stem cells states colorectal cancer resections, and the expression of the markers *CDKN1A/P21, CDKN1B/P27, CDKN1C/P57, LRIG1 and MUC2*. **(b)** Heatmap displaying the various ECM molecular signatures against the clusters that were generated using the whole transcriptome. **(c)** Heatmap displaying the various ECM molecular signatures against the clusters that were generated using ECM genes only. **(d)** Heatmap displaying the various ECM molecular signatures against the clusters that were generated using MPs.

Figure 5 Suppl 1

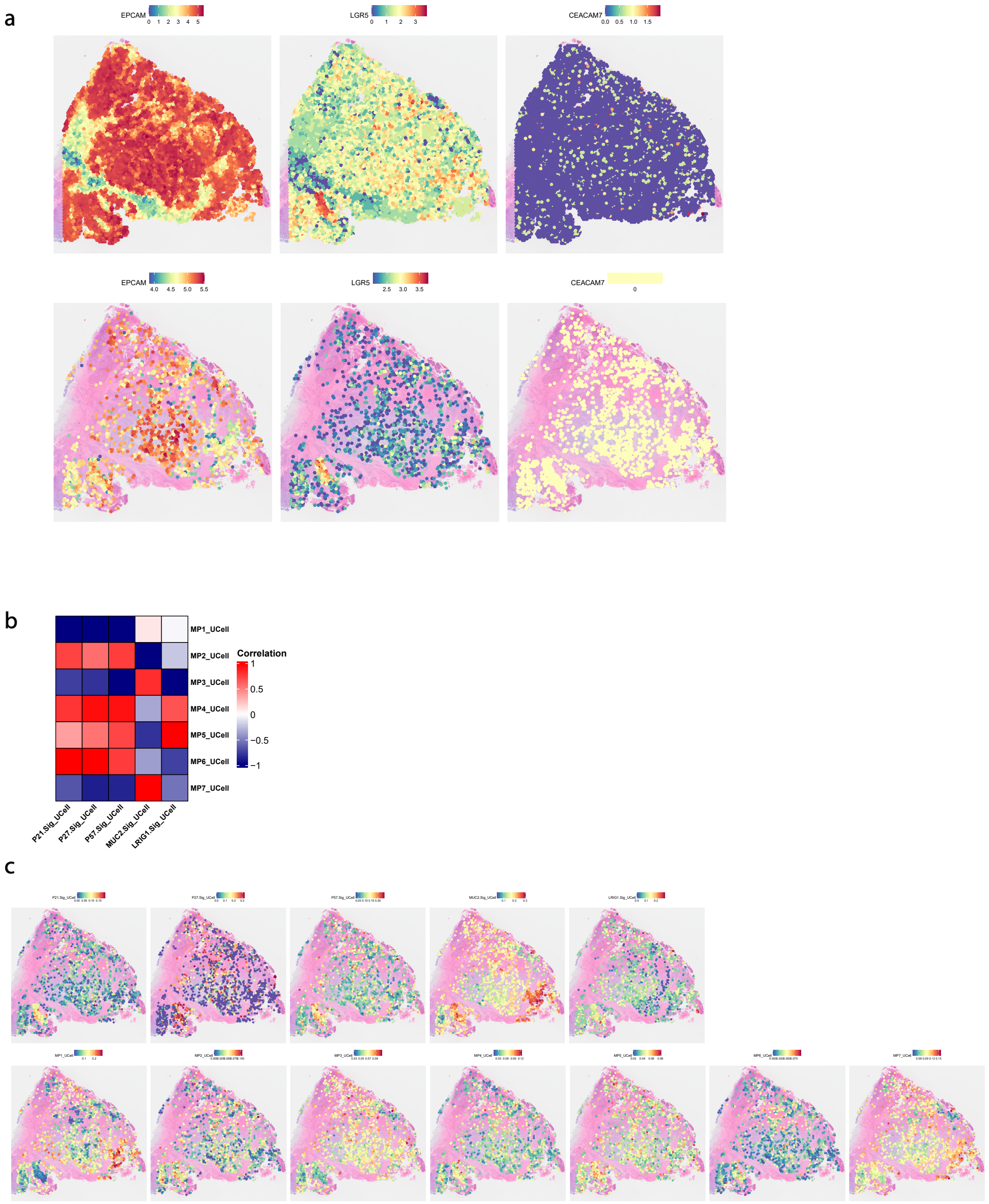


**Figure 5. S1.** **(a)** Spatial DimPlots showing the expression of EPCAM, LGR5 and CEACAM7 across every region of interest (n = 9080) in the CRC tissue section. This is followed by Spatial DimPlots showing the expression of EPCAM, LGR5 and CEACAM7 across the subsetted regions of interest (n = 1658). **(b)** Heatmap showing the correlation between the ECM signature scores and the MP signature scores across the subsetted stem-like regions of interest in the CRC tissue section. **(c)** Spatial FeaturePlots demonstrating the expression of each of the CSC ECM based molecular signatures, and the signatures driving the MPs for CSCs across the CRC tissue section.

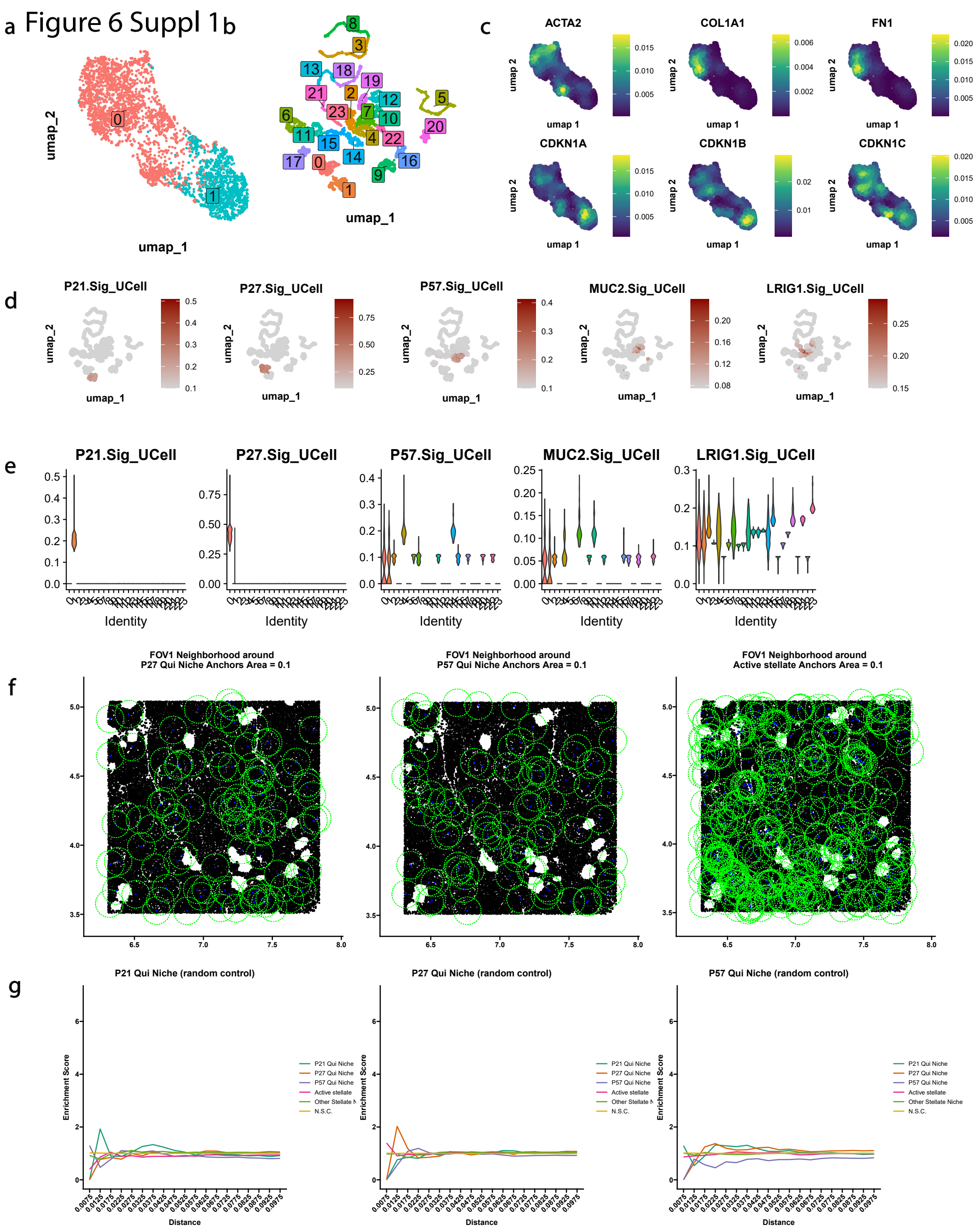


**Figure 6. S1.** (**a**) UMAP showing two distinct clusters of pancreatic stellate cells. (**b**) UMAP showing twenty-four clusters of pancreatic stellate cells based on clustering using the healthy intestinal ECM based molecular signatures. (**c**) Density plots exhibiting the expression of activated stellate cell markers; ACTA2, COL1A1 and FN1, and quiscent cell markers; CDKN1A/P21, CDKN1B/P27 and CDKN1C/P57. (**d**) Feature plots displaying the expression of the ECM signatures for various stem cell states overlaid on cells in the UMAP from (**b**). (**e**) Violin plots that show the expression of each healthy intestinal ECM based molecular signature in each of the stellate cell clusters. (**f**) Scatter plots of the location of each cell type within the pancreas tissue field of view. Anchor cells are coloured in blue with the maximal radial distance shown in the green dotted circle surrounding each anchor cell. (**g**) Neighborhood analysis plots that shows the enrichment of each cell type above the background frequency surrounding each anchor cell type at increasing distances up to 0.1 mm.
